# A dietary pan-amino acid dropout screen *in vivo* reveals a critical role for histidine in T-ALL

**DOI:** 10.64898/2025.12.21.694897

**Authors:** Komal Mandleywala, Simona Ulrich, Victoria da Silva-Diz, Puneet Sharma, Christopher Thai, Oekyung Kim, Maya Aleksandrova, Gwendolyn Chung, Cristian Eggers, Dieter Lütjohann, Tanaya Kulkarni, Amartya Singh, M. Elena Díaz-Rubio, Michael Wierer, Sebastian A. Leidel, Xiaoyang Su, Eileen P. White, Joshua D. Rabinowitz, Raphael J. Morscher, Daniel Herranz

**Affiliations:** Rutgers Cancer Institute, Rutgers University, New Brunswick, NJ, 08901, USA; Pediatric Cancer Metabolism Laboratory, Children’s Research Center, University of Zurich, 8008, Zurich, Switzerland; Division of Oncology, University Children’s Hospital Zurich and Children’s Research Center, University of Zurich, 8008, Zurich, Switzerland; Department of Biology, Institute of Biochemistry, ETH Zurich, 8093 Zurich, Switzerland; NCCR RNA and Disease Technology Platform, Bern, Switzerland; Center for Systems and Computational Biology, Rutgers Cancer Institute, Rutgers University, New Brunswick, NJ, 08901, USA. 20892, USA; Department of Chemistry, Biochemistry and Pharmaceutical Sciences, University of Bern, 3012 Bern, Switzerland; Institute of Clinical Chemistry and Clinical Pharmacology, University Hospital Bonn, 53127 Bonn, Germany; Department of Medicine, Rutgers Robert Wood Johnson Medical School, Rutgers University, New Brunswick, NJ, 08901, USA; Proteomics Research Infrastructure, Panum Institute, University of Copenhagen, Copenhagen, Denmark; Ludwig Institute for Cancer Research, Princeton Branch, Princeton University, Princeton, NJ 08544, USA; Department of Molecular Biology and Biochemistry, Rutgers University, Piscataway, NJ 08901, USA; Department of Chemistry, Princeton University, Princeton, NJ 08544, USA; Department of Pharmacology, Rutgers Robert Wood Johnson Medical School, Rutgers University, Piscataway, NJ, 08854, USA; Department of Pediatrics, Rutgers Robert Wood Johnson Medical School, Rutgers University, New Brunswick, NJ, 08901, USA

**Author notes:** **Corresponding authors:** Raphael J. Morscher, Group Leader - Pediatric Cancer Metabolism Laboratory, Children’s Research Center, August-Forel-Strasse 51, 8008, Zürich, Switzerland, Daniel Herranz, Associate Professor of Pharmacology and Pediatrics, Robert Wood Johnson Medical School, Rutgers Cancer Institute, Rutgers, The State University of New Jersey, 195 Little Albany Street, Office Room 3037, Lab Room 3026, New Brunswick, NJ, 08901. Equal contribution (Komal Mandleywala, Simona Ulrich, Victoria da Silva-Diz).

**Keywords:** T-cell acute lymphoblastic leukemia, T-ALL, Dietary interventions, Histidine, Cholesterol

## Abstract

Dietary interventions show therapeutic potential in cancer, but systematic comparisons are lacking. We performed a dietary pan-amino acid dropout screen in an orthotopic model of NOTCH1-driven T-cell acute lymphoblastic leukemia and identified histidine depletion as uniquely antileukemic. Histidine-restricted diets extended survival of leukemic mice in a dose-dependent manner, while remaining well-tolerated. Mechanistically, multiomic profiling revealed that histidine deprivation-induced ribosome stalling activates GCN2 to suppress cholesterol biosynthesis pathways critical for leukemic proliferation. Dietary cholesterol supplementation partially reverted the antileukemic effects of histidine restriction *in vivo*. These findings couple histidine levels and translational control to cholesterol metabolism, which can be therapeutically exploited for cancer treatment. Our results suggest that defined dietary amino acid restrictions may expose broader therapeutic opportunities in diseases beyond cancer.

---

T-cell acute lymphoblastic leukemia (T-ALL) is an aggressive NOTCH1-driven hematological malignancy affecting both children and adults^1,2^. Despite intensified chemotherapy protocols that have improved outcomes, 20-50% of patients still relapse with dismal prognosis^3^. Even if cured, survivors face significant therapy-induced toxicities^3^, underscoring the need to identify novel and safer treatments. Unlike recent advances in immunotherapy for B-ALL treatment using CAR T-cells or antibodies, these approaches are more challenging in T-ALL^4–6^. In contrast, targeting metabolism has long been a cornerstone of leukemia therapy, as exemplified by the clinical use of methotrexate, 6-mercaptopurine or L-asparaginase^7^, highlighting the value of further exploring metabolic vulnerabilities in leukemia. Building on this, we recently identified additional metabolic interventions with antileukemic effects by targeting either serine hydroxymethyltransferase, ATP citrate-lyase or mitochondrial respiration^8–10^. Moreover, modulation of organismal metabolism through dietary interventions has gained significant attention in recent years based on promising therapeutic effects in preclinical cancer models^11–13^, and are currently being evaluated in clinical trials (e.g. NCT05300048 or NCT01535911). Still, most studies have focused on single dietary interventions and have not assessed their effects in primary orthotopic systems. Given the amenability of T-ALL to metabolic targeting, we hypothesized that modulating nutrient composition in the diet might lead to effective antileukemic effects. To test this hypothesis, we leveraged the unique strengths of our NOTCH1-induced leukemia mouse model (primary, orthotopic, immune-competent and high-throughput) and performed dietary pan-amino acid dropout screens *in vivo*. Unexpectedly, these unbiased experiments revealed the strongest intrinsic antileukemic effect for diets lacking histidine.

## Dietary amino-acid dropout screens *in vivo* identify a critical role for histidine in T-ALL progression

To test the antileukemic effects of dietary interventions, we induced T-ALL by transducing an oncogenic form of NOTCH1 in murine bone marrow progenitor cells, followed by transplantation into lethally irradiated recipients^14^ (**Fig. 1a**). Subsequently, leukemias were transplanted into secondary mice randomly assigned to feeding either with a control diet, or diets lacking each of the 20 amino acids while maintaining the total protein composition constant (**Fig. 1a**). Interestingly, this experiment revealed three broader effects: First, removal of most amino acids did not impact T-ALL progression (**Fig. 1b**). These included diets without asparagine, tyrosine, proline, glutamine, isoleucine, glutamate, cysteine, arginine, alanine, serine or glycine (**Fig. 1b**). A second group − diets without tryptophan, threonine, phenylalanine, lysine or aspartate − showed significant, but delayed antileukemic effects (**Fig. 1c**). These diets led to a bimodal effect on survival: initial mice succumbed with similar kinetics to controls, but the remaining mice showed a significant extension in survival (**Fig. 1c**). The third group of diets was the most interesting one, showing strong intrinsic antileukemic effects from the outset (**Fig. 1d**). These included diets without valine, leucine, methionine and histidine, leading to significant survival extension not driven by decreased food intake (**Fig. 1d and Extended Data Fig. 1a**). Importantly, valine restriction has been previously shown to result in antileukemic effects^15^, validating our results. However, both valine and leucine restriction were severely toxic, as expected from the lack of essential amino acids, and mice lost significant weight in the first weeks (**Extended Data Fig. 1b**), forcing us to switch them back to control diets even before they died from leukemia (**Fig. 1d**). These results suggest that targeting these amino acids in patients might be clinically challenging. Conversely, diets without methionine and histidine were better tolerated (**Extended Data Fig. 1b**). Methionine restriction has previously shown therapeutic effects in multiple models of solid cancer^16,17^ and acute myeloid leukemia^18^. On the other hand, dietary histidine restriction led to a 5-fold decrease in serum histidine levels (**Extended Data Fig. 1c**) and showed the strongest therapeutic effects of them all (**Fig. 1d**). Moreover, the role of dietary histidine in cancer treatment is largely unknown. Therefore, we focused our efforts on dissecting the effects of dietary histidine in T-ALL progression.

**Fig. 1.**
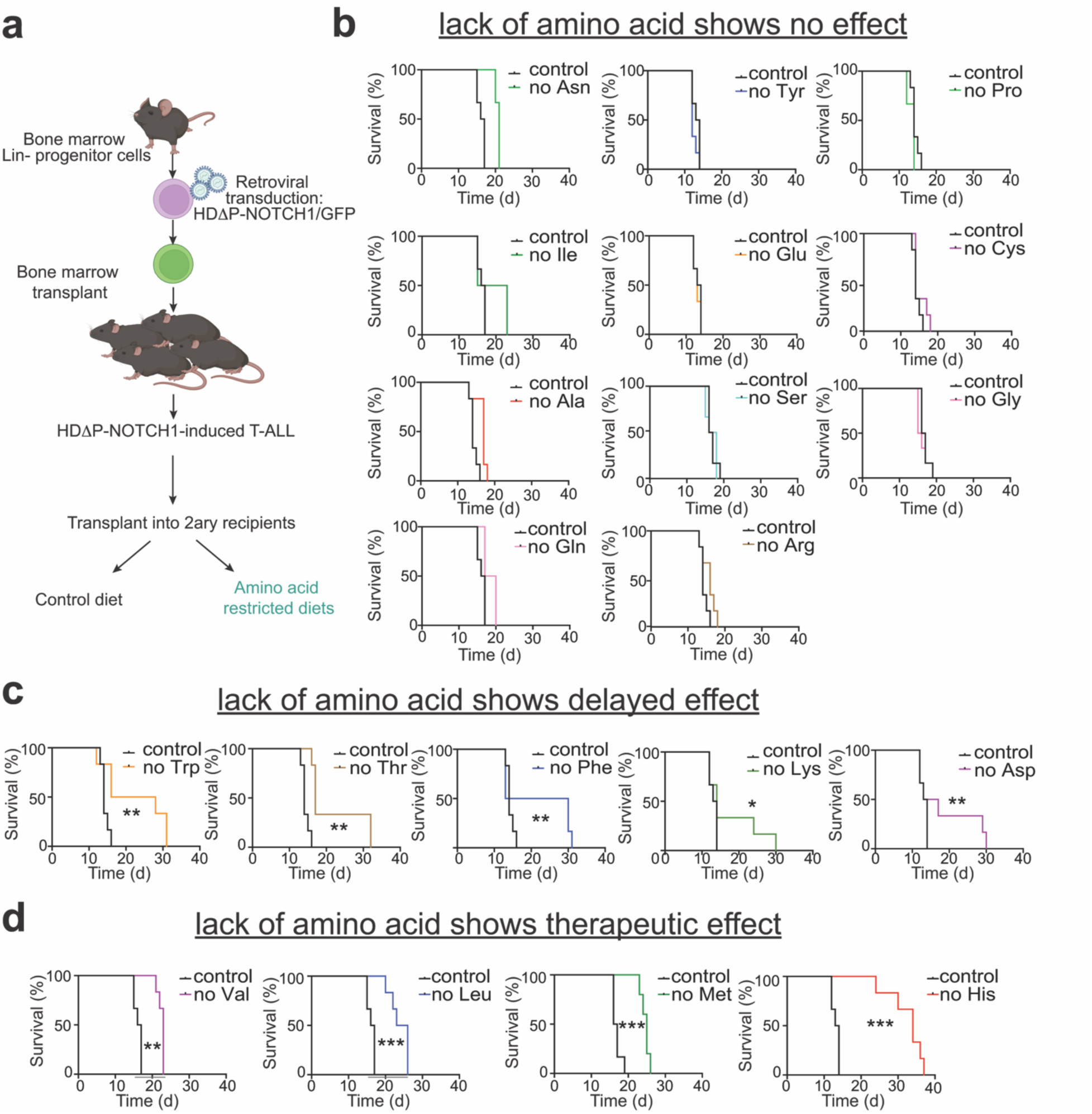
Dietary amino-acid dropout screens in T-ALL *in vivo* uncover significant therapeutic effects for histidine restriction. **a**, Schematic of NOTCH1-induced T-ALL generation and workflow for survival analyses upon treatment of leukemic mice with diets lacking different amino acids. **b**−**d**, Kaplan-Meier survival curves of mice harboring NOTCH1-induced T-ALL and fed with control diet or with diets lacking one amino acid that either showed no effect on T-ALL progression (**b**); showed delayed but significant therapeutic effects (**c**); or showed significant intrinsic antileukemic effects (**d**). The colored bars below the X axis in “no Val” and “no Leu” survivals indicate diet discontinuation and replacement by control diet, due to mouse toxicity. For the rest of the experiments, amino acid-deficient diets were given continuously from day 2 post leukemic cell transplantation up until mouse euthanasia. \**P* < 0.05; \*\**P* < 0.01; \*\*\**P* < 0.005 calculated with log-rank test (n = 6 per group).

## Histidine shows dose-dependent effects and histidine restriction is therapeutic in human T-ALL patient-derived xenografts (PDXs) *in vivo*

While dietary histidine absence showed strong antileukemic effects, complete histidine elimination from patient diets would likely be a considerable challenge in practice. We thus next tested whether reducing (but not eliminating) dietary histidine would show therapeutic effects in mice with primary NOTCH1-driven leukemias. We used diets with half (50% His), one fourth (25% His) or one tenth (10% His) of normal histidine levels. Importantly, we observed dose-dependent antileukemic effects upon histidine reduction *in vivo* (**Fig. 2a**), suggesting that tailored diets with reduced histidine may have antileukemic effects in patients. We also tested whether increasing dietary histidine would produce the opposite effects. Indeed, leukemic mice fed diets containing 10-fold normal histidine levels showed accelerated disease kinetics (**Fig. 2b**), demonstrating that dietary histidine has dose-dependent effects in T-ALL progression *in vivo*. Finally, we tested dietary histidine restriction in human T-ALL PDXs in immunodeficient mice. Of note, we observed significant survival extension in mice transplanted with T-ALL PDXs from two clinically distinct molecular subgroups (PDTALL-10^10^, TLX3 subgroup; PDTALL-13^19^, TAL-LMO subgroup; **Fig. 2c**), and these PDXs were either wild-type or mutated for *NOTCH1* and *PTEN* (**Fig. 2c**). These results suggest that histidine restriction has therapeutic effects in T-ALL irrespective of mutational drivers. Moreover, they support that the antileukemic effects are at least partly tumor cell-intrinsic, as histidine restriction increased survival even in PDX-harboring immunodeficient mice.

**Fig. 2.**
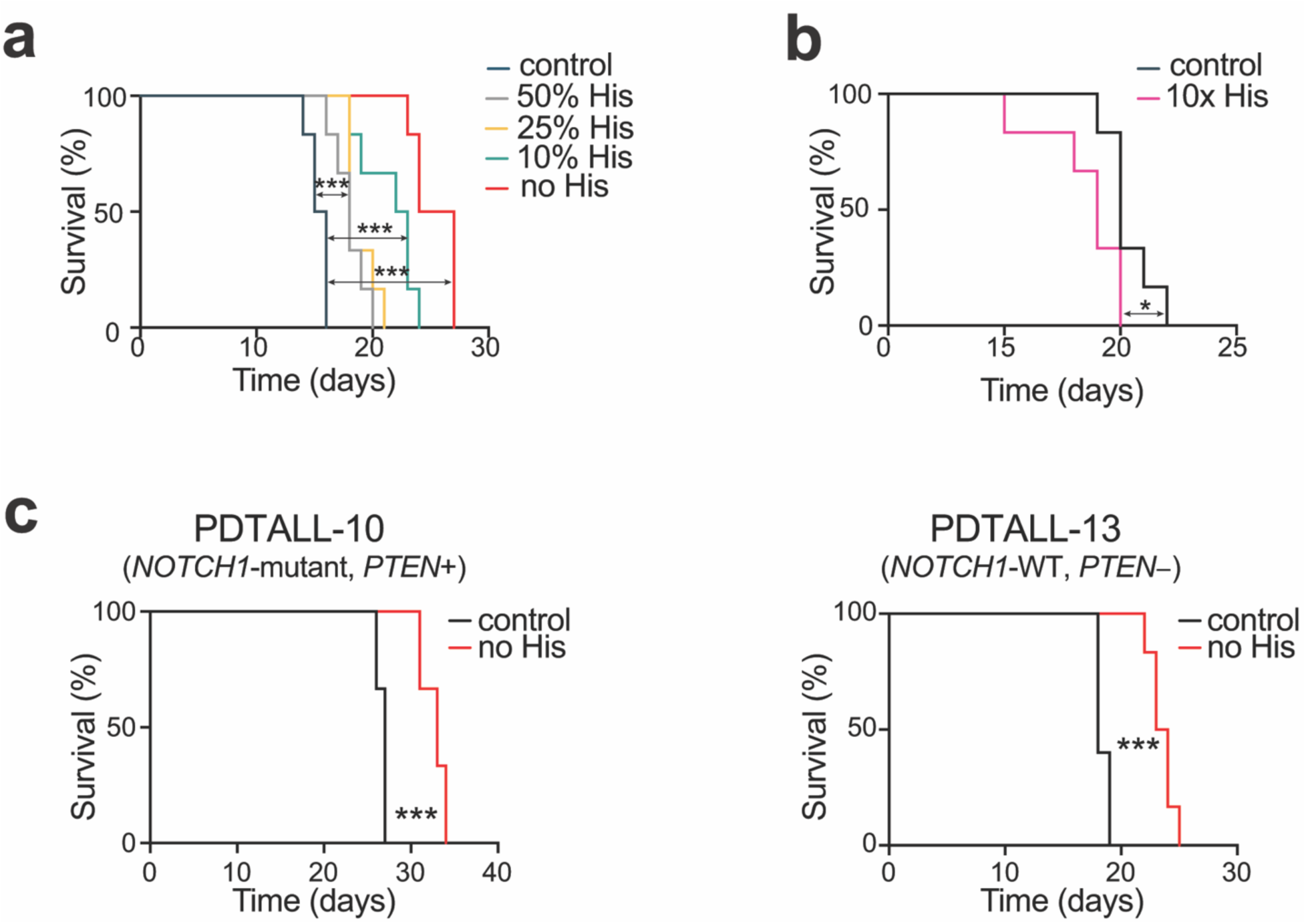
Dose-dependent effects of dietary histidine in T-ALL progression. **a**, Kaplan-Meier survival curves of mice harboring NOTCH1-induced T-ALL and fed with control diet or with diets with decreasing amount of histidine (50%, 25%, 10% or 0% of the levels in control diet). **b**, Kaplan-Meier survival curves of mice harboring NOTCH1-induced T-ALL and fed with control diet or with a diet having a 10-fold excess in histidine compared to normal histidine levels. **c**, Kaplan-Meier survival curve of immunodeficient mice harboring a *NOTCH1*-mutant and PTEN-positive human T-ALL patient-derived xenograft (PDX; left) or a *NOTCH1*-wild type and PTEN-negative human T-ALL PDX (right), fed with control diet or with a diet without histidine. \**P* < 0.05; \*\*\**P* < 0.005 calculated with log-rank test (n = 6 per group).

## Dietary histidine is critical for normal T-cell development

Since T-ALL therapies often affect normal T-cell development concomitantly, we next tested whether dietary histidine restriction in healthy mice impairs thymocyte development (**Fig. 3a**). Notably, mice fed histidine-free diets for only 2 weeks showed drastic reductions in thymus weight and cellularity (**Fig. 3b,c**). Detailed immunophenotypic analyses showed significant reductions in all thymocyte subsets, including CD4/CD8 double-negative (DN: DN1-DN4), double positive (DP), and CD4 or CD8 single-positive (SP) thymocytes (**Fig. 3d,e**). This phenotype was due to an early developmental block at the DN stage, with increased proportions of total DN thymocytes, also evident at the earliest stage of DN1 cells (**Fig. 3d,f**). Healthy mice fed histidine-free diets long-term (over a month) progressively lost weight (**Fig. 3g**). Bloodwork revealed a significant reduction in total white blood cell counts (**Fig. 3h**), selectively affecting lymphocytes, monocytes, eosinophils and basophils, while neutrophils remained unaffected (**Fig. 3h**). Conversely, red blood cells and platelets showed minimal changes (**Fig. 3h**). These results demonstrate that dietary histidine is critical for both T-ALL progression and normal T-cell development. The limited hematological toxicity across other blood cell populations supports further exploration in clinical translation.

**Fig. 3.**
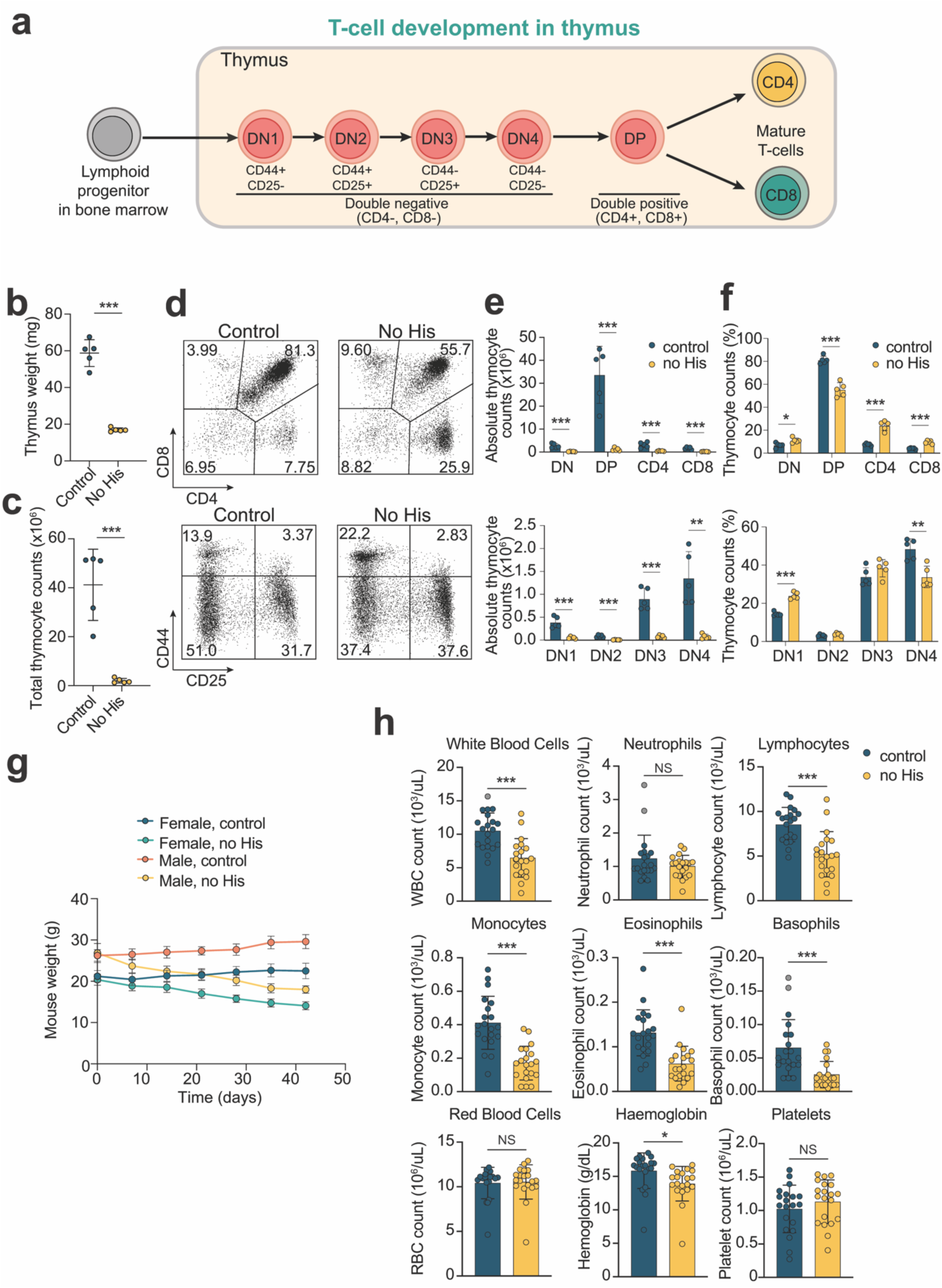
Effects of histidine-deficient diet on normal T-cell development and peripheral blood counts. **a**, Schematic of normal T-cell development in the thymus. **b,** Thymus weight in normal wild-type mice fed a control diet or a diet without histidine for 2 weeks. **c**, Total thymocyte count in normal wild-type mice fed a control diet or a diet without histidine for 2 weeks. **d**−**f**, Representative plots (**d**), absolute quantification (**e**) or relative quantification (**f**) of CD4−CD8− (double negative: DN), CD4+CD8+ (double positive: DP), CD4+CD8− (CD4), CD4−CD8+ (CD8), CD44+CD25− (double negative 1: DN1), CD44+CD25+ (double negative 2: DN2), CD44−CD25+ (double negative 3: DN3), and CD25−CD44− (double negative 4: DN4) populations in the thymus of normal wild-type mice fed a control diet or a diet without histidine for 2 weeks. Data are mean ± s.e.m (n=5 per group). **g**, Change in mouse weight upon long-term >1-month feeding with control diet or diet without histidine. Data are mean ± sem (n=10 per group). **h**, Changes in bloodwork parameters in mice upon long-term >1-month feeding with control diet or diet without histidine. Data are mean ± sem (n=20 per group). NS = not significant; \**P* < 0.05; \*\**P* < 0.01; \*\*\**P* < 0.005 using two-tailed Student’s *t*-test.

## Histidine restriction results in downregulation of cholesterol-related pathways

To dissect the mechanistic consequences of histidine restriction, we explored its effects in human T-ALL cell lines *in vitro*. As expected, histidine restriction led to a dose-dependent impairment in proliferation of Jurkat cells (**Fig. 4a,b**), and across a broader panel of T-ALL cell lines (**Extended Data Fig. 2a**−**d**). These effects combined cytotoxic and cytostatic mechanisms, with progressive increases in Annexin V staining (**Fig. 4c**) and cell cycle arrest in G0 (**Fig. 4d**) as histidine decreased in the culture media. We next performed global metabolomics to assess how histidine restriction affects T-ALL metabolism. As anticipated, histidine restriction led to a dose-dependent decrease in intracellular histidine (**Fig. 4e**) with prominent accumulation of glutamine and serine (**Fig. 4e**). We also observed significant pyruvate accumulation with concomitant depletion in TCA cycle intermediates, nucleotide intermediates and metabolites with reducing potential, such as NADH and glutathione (**Extended Data Fig. 2e**−**g**). However, tracing experiments with fully labeled U-^13^C-labeled histidine showed negligible TCA and nucleotide intermediate labeling both after short-term culture or after labeling for over one week (**Extended Data Figs. 3 and 4**). These results suggest that histidine catabolism is not a meaningful nutrient source.

**Fig. 4.**
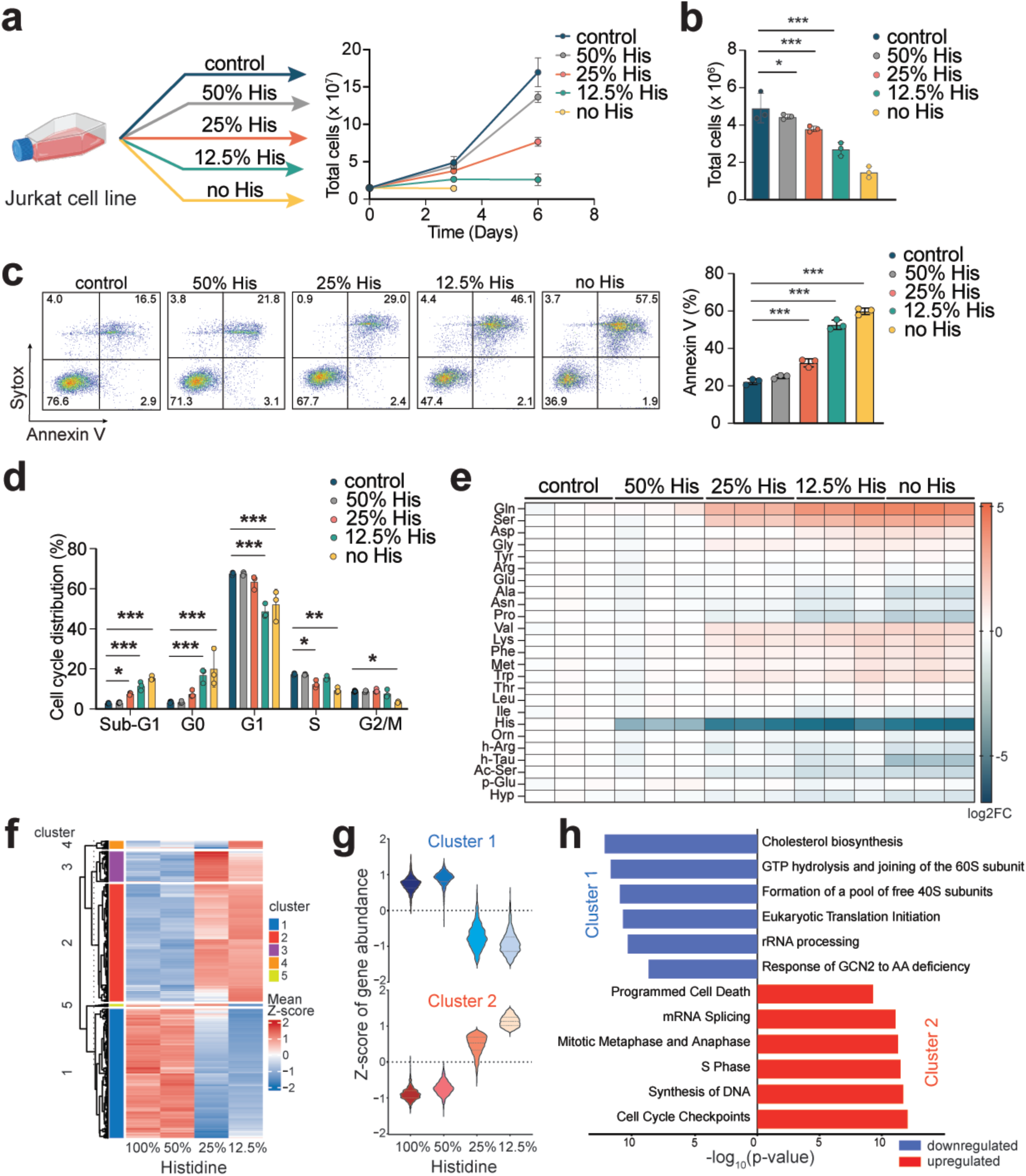
Histidine restriction *in vitro* leads to antileukemic effects and downregulation of cholesterol transcriptional signatures. **a**, Schematic (left) and proliferation curve (right) of Jurkat T-ALL cells grown in normal RPMI media, RPMI with 50%, 25% or 12.5% levels of histidine, or RPMI without histidine. **b**, Quantification of changes in proliferation at day 6. Data are mean ± sem (n=3). **c**, Representative flow cytometry plots of annexin V and Sytox staining (left) and quantification (right) from Jurkat triplicates grown for 72 h in normal RPMI media or RPMI with decreasing histidine levels. Numbers in quadrants indicate percentage of cells. Data are mean ± sem (n=3). **d**, FACS- based quantification of cell cycle analysis of Jurkat triplicates grown for 72 h in normal RPMI media or RPMI with decreasing histidine levels. Data are mean ± sem (n=3). *P* values in b−d were calculated using two-way ANOVA for multiple comparisons; \**P* < 0.05; \*\**P* < 0.01; \*\*\**P* < 0.005. **e**, Heatmap showing differential intracellular amino acid abundances (log_2_) in Jurkat triplicates grown for 72 h in RPMI with decreasing histidine levels, relative to RPMI-control. **f**, Heatmap depicting five gene expression clusters in response to decreasing histidine levels (100–12.5%), with colors indicating mean Z-scores (n=3). **g**, Selected clusters of genes based on their continuous decreasing/increasing trends across histidine levels. **h**, Pathway analyses using gProfiler2 with the list of significantly down/upregulated genes (g:SCS method; *P* < 0.01).

In order to better understand the potential mechanisms underlying histidine restriction, we performed gene expression profiling by RNA-seq in Jurkat cells cultured with decreasing histidine concentrations. We observed two clearly differentiated groups: cells cultured in 100% or 50% of histidine clustered together, whereas cells grown in 25% or 12.5% histidine showed distinct transcriptional signatures (**Fig. 4f,g**). Surprisingly, pathway analyses using gProfiler2^20^ revealed striking downregulation of cholesterol-related processes upon histidine restriction (**Fig. 4h**). Similar results were obtained in the independent human T-ALL cell line DND41 (**Extended Data Fig. 5a,b**), indicating this is a conserved phenotype upon histidine limitation. Since gene sets in pathway analyses sometimes encompass loosely related genes, we validated cholesterol’s relevance by generating a cholesterol score, using only genes directly related to cholesterol synthesis. These analyses, analogous to the OxPhos score^21^, confirmed the strong downregulation in cholesterol biosynthesis in both cell lines (**Extended Data Fig. 5c**). While the link between histidine and cholesterol was completely unexpected, thorough analyses of decades-old literature revealed that rats fed diets with excess in histidine were previously found to develop hypercholesterolemia^22,23^. Finally, we wondered whether restriction of any essential amino acid, and not only histidine, would lead to downregulation of cholesterol metabolism. Indeed, Jurkat cells grown in 25% levels of most essential amino acids showed reduced proliferation and a strong downregulation of the transcriptional cholesterol score (**Extended Data Fig. 5d**−**e**). These findings suggest that the cholesterol pathway downregulation might indeed be relevant for the therapeutic effects of essential amino acid restrictions in T-ALL and, more broadly, support an underappreciated relevance of the histidine-cholesterol axis.

## Histidine restriction rewires protein translation

Given the lack of contributions from histidine degradation to downstream metabolites, we next investigated the impact of limited histidine supply as a substrate for translation. Histidine codon frequency in proteins did not predict global pathway regulation on proteomics (**Extended Data Fig. 6a,b**), despite proteins with a high histidine content being significantly downregulated across the proteome (**Extended Data Fig. 6c**). Specifically, we found no correlation between histidine content and the expression of cholesterol biosynthesis enzymes (**Extended Data Fig. 6d**), pointing towards a more complex sensing response. Across species, highly conserved mechanisms regulate protein production rates based on amino acid availability^24^. In cancer, this also allows transformed cells to tightly coordinate biosynthetic demands with nutrient scarce environments. Ribosome profiling (Ribo-seq) has served as a valuable tool to dissect the consequences of amino acid limitation on translation dynamics at codon resolution by sequencing mRNA fragments protected by ribosomes at the moment of lysis^25^ (**Fig. 5a and Extended Data Fig. 7**). When assessing translation dynamics by Ribo-seq upon histidine restriction, we observed a striking increase of ribosome density (i.e. stalling) at both histidine codons (CAC and CAU) in the A-site, indicating that these codons are decoded slowly presumably due to uncharged His-tRNAs (**Fig. 5b**). Interestingly, inspection of ribosome density around these CAC and CAU codons revealed an additional upstream stalling peak (**Fig. 5c**). The 10-codon distance between the major and minor peaks corresponds to the spacing between the A-sites of a leading stalled ribosome and a trailing ribosome that has collided with it^26^. No such peaks were observed for any other codons (e.g. AAA, GAA, AAG and GAG) (**Fig. 5c**).

**Fig. 5.**
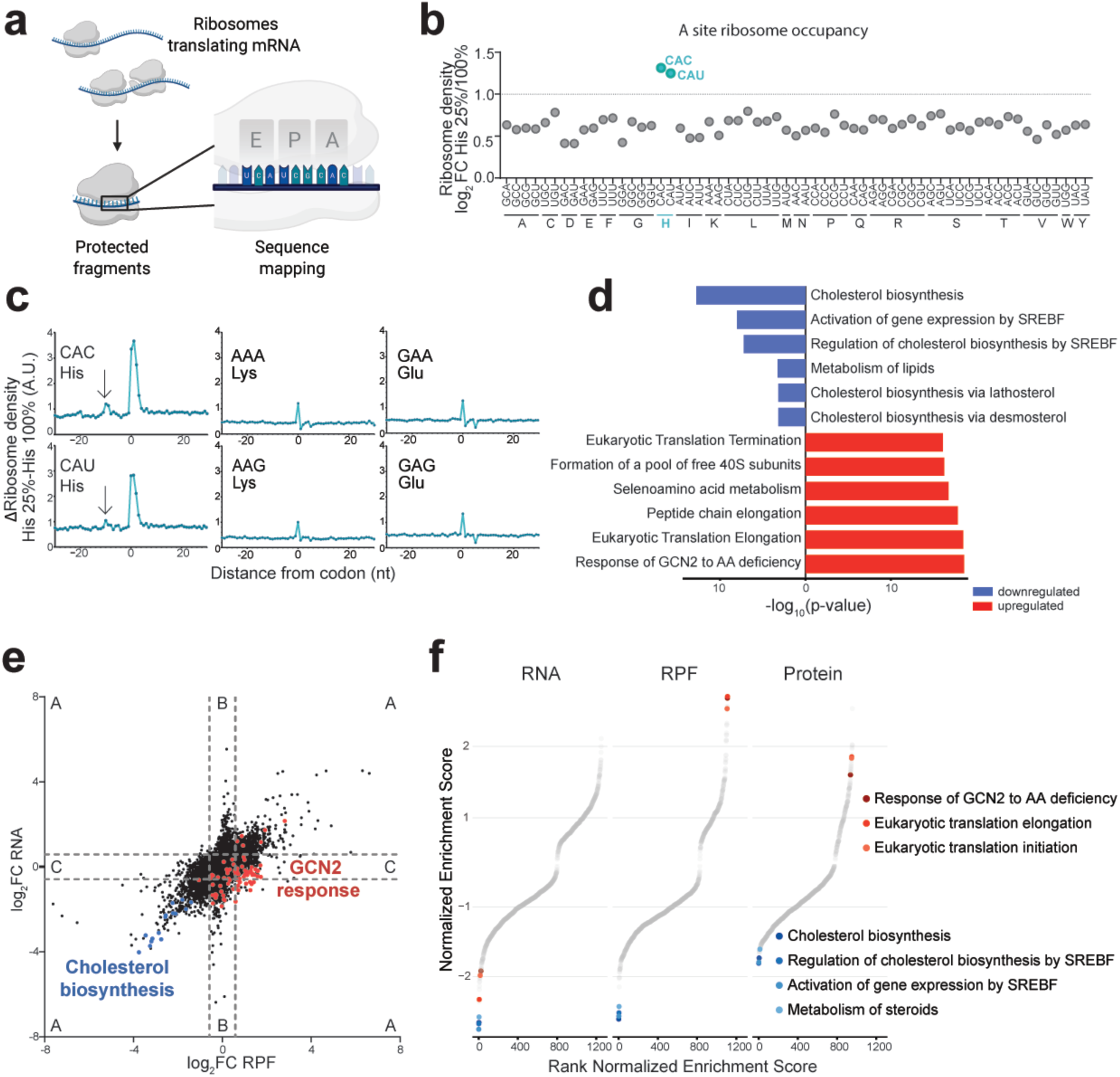
Histidine restriction rewires translation and affects cholesterol biosynthesis. **a**, Schematic of ribosome profiling with protected fragments being sequenced and mapped to the ribosome A-site. **b**, Relative ribosome density across all codons in Jurkat cells grown in 25% His, as compared to control RPMI conditions. Pronounced stalling is observed at His-coding codons (CAC and CAU) (n=3). **c**, A-site ribosome density profiles around given codon identities (at position 0) reveal specific stalling at histidine codons compared to representative controls lysine (AAA, AAG) and glutamate (GAA, GAG). The density peaks at -10 codons (arrows) in CAC and CAU profiles indicate collided ribosomes (n=3). **d**, Most significantly regulated pathways (Reactome) in gene set enrichment analysis (GSEA) at the translational level (Ribo-seq) in Jurkat cells grown in 25% His. **e**, Relation of RNA abundance and ribosome occupancy in matched samples highlights the respective regulation of expression at the different levels (A regulated on transcription and translation level; B transcription only; C translation only). **f**, Corresponding GSEA analysis across omics datasets comparing Jurkat cells grown in 25% His vs control. All depicted Reactome pathways are ranked by normalized enrichment score. Among the most down- and upregulated sets are ‘Cholesterol biosynthesis’ and ‘Response of GCN2 to amino acid deficiency’, respectively.

In the T-ALL context, histidine restriction therefore induces ribosome stalling, rather than merely suppressing translation initiation^27^. Such stalling with ribosome collisions has recently been shown to be essential for GCN2 kinase activation^28^, which we confirmed by western blot analyses (**Extended Data Fig. 8a**). Concomitantly, we observed a selective activation of the co-translational branch of the unfolded protein response (PERK)^29^ (**Extended Data Fig. 8a**). Both pathways converge in mammalian cells on the conserved integrated stress response (ISR) as a coordinated response to amino acid deprivation or proteostasis defects^30^, regulating translation and ATF4 activation (**Extended Data Fig. 8a**). In contrast, the amino acid sensor mTOR^31^ showed a translation-promoting activation (**Extended Data Fig. 8b**), validating its inability to sense histidine levels^32^. Net reduced protein synthesis rates were evident upon puromycin incorporation rates (**Extended Data Fig. 9**). Consistently, unbiased pathway enrichment analysis of Ribo-seq upon histidine limitation highlighted GCN2 activation signatures and pathways related to translation (**Fig. 5d−f**). This upregulation at the ribosome and protein level was not seen at the mRNA level, implicating a post-transcriptional response (**Fig. 5e−f**). Surprisingly, on the opposite spectrum, cholesterol biosynthesis was again the most suppressed pathway also at the translational and proteomic level, highlighting its consistent downregulation across different omics (**Fig. 5e,f and Extended Data Fig. 8c**).

## GCN2 activation upon histidine depletion suppresses cholesterol biosynthesis

Cholesterol biosynthesis is a tightly regulated multi-enzyme pathway coordinated with uptake and efflux to meet cellular demands (**Fig. 6a**) and its homeostasis is primarily orchestrated by the master regulator sterol regulatory element-binding protein 2 (SREBF2)^33,34^. While cholesterol biosynthesis was globally downregulated, it is important to note that, at the protein level, this prominently included SREBF2 (**Fig. 6a,b**), which might be mediating the observed phenotype. These findings were confirmed by western blot analyses upon acute histidine restriction showing reduced levels of SREBF2, HMGCR and HMGCS1, together with the phosphorylation of GCN2 (**Fig. 6c**) and PERK, and accumulation of ATF4 (**Extended Data Fig. 8a**). Interestingly, blocking ISR activation upon histidine restriction by pharmacologic inhibition of GCN2 phosphorylation restored both the expression of cholesterol biosynthesis enzymes and leukemic cell proliferation (**Fig. 6d and Extended Data Fig. 10a**). Conversely, pharmacologic activation of GCN2 was sufficient to induce a strong downregulation of HMGCS1 in histidine-rich media (**Fig. 6e**), while histidine depletion conferred dose-dependent resistance against its growth inhibition (**Extended Data Fig. 10b**). Overall, these results highlight a key role of the GCN2-mediated stress response signaling upon histidine deprivation to induce downregulation of cholesterol biosynthesis.

**Fig. 6.**
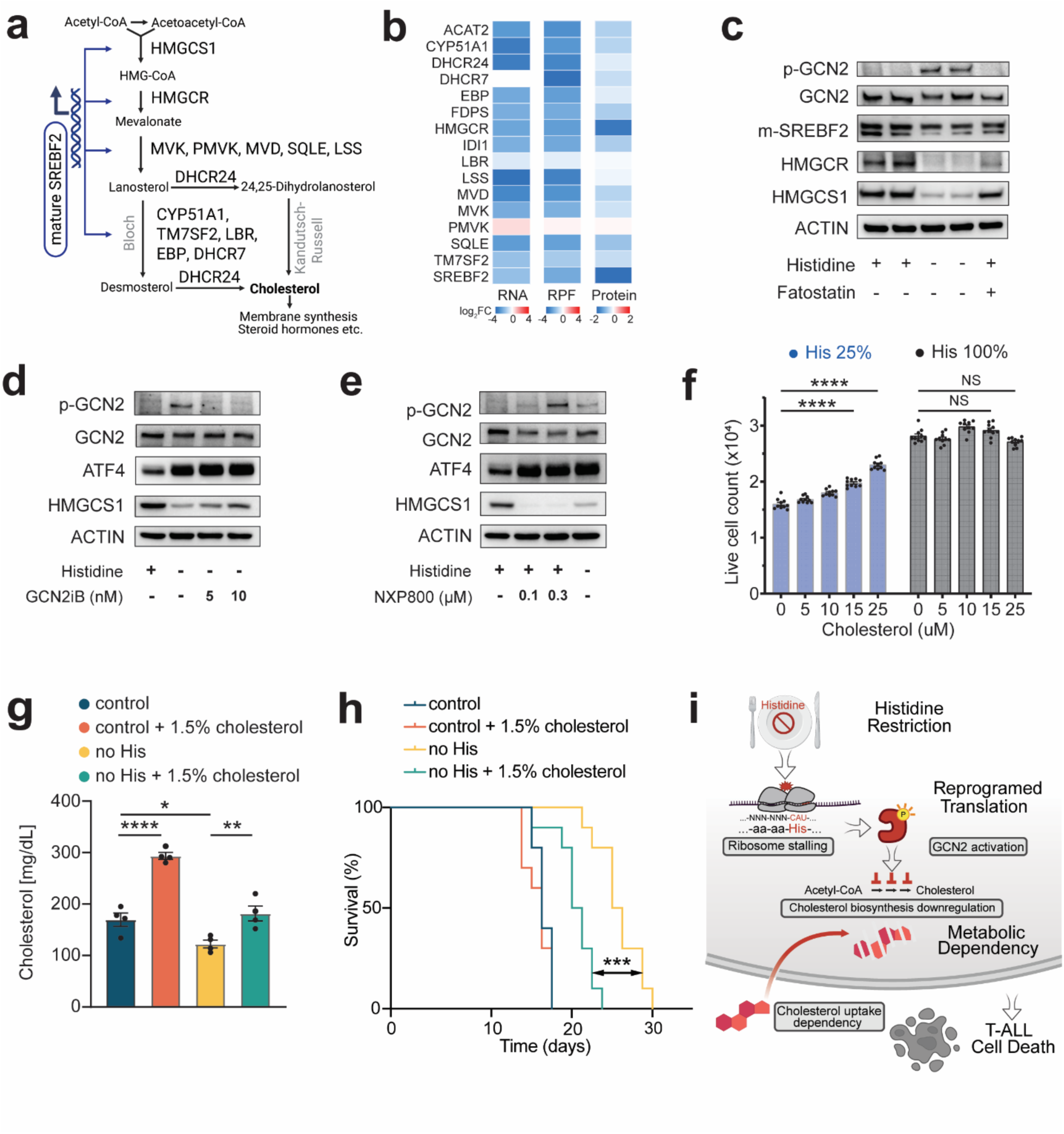
Histidine restriction rewires translation and induces cholesterol-dependency. **a**, Schematic of cholesterol biosynthesis pathway. **b**, Heatmap showing Log_2_FC downregulation of cholesterol-related enzymes and the primary regulator SREBF2 across RNA-seq, Ribo-seq and proteomics in Jurkat cells. **c**, Western blot analyses of GCN2 pathway members (phospho-GCN2, total GCN2) and respective regulation of the cholesterol pathway (SREBF2, HMGCR, HMGCS1) in Jurkat cells grown in 25% His for 72 h and treated with either DMSO or fatostatin, an inhibitor of SREBF2 activation. **d,** Suppression of GCN2 activation by GCN2iB partially rescues cholesterol biosynthesis enzymes in Jurkat cells upon His restriction (25%), as shown by western blot analyses. **e**, Reversely, pharmacological activation of GCN2 by NXP800 leads to HMGCS1 downregulation independently of His levels. **f**, The proliferation of Jurkat cells grown in under 25% His shows a dose-dependent rescue by cholesterol supplementation when grown in delipidated fetal bovine serum. Data are mean ± sem (n=8). **g,** Serum cholesterol levels in mice harboring NOTCH1-induced T-ALL and fed with control or histidine-free diet, highlighting cholesterol decrease in mouse serum upon histidine depletion. The diets are enriched (or not) with 1.5% cholesterol. Data are mean ± sem (n=4). **h,** Kaplan-Meier survival curves of mice harboring NOTCH1-induced T-ALL and fed with control diet or diets without histidine and supplemented (or not) with 1.5% cholesterol. For f and g, *P* value calculated using two-way ANOVA for multiple comparisons; NS = not significant; \**P<0.05,* \*\**P<0.01,* \*\*\*\**P* < 0.0001. For h, Kaplan-Meier survival \*\*\**P* < 0.005 calculated with log-rank test (n = 10 per group). **i,** Overview schematic highlighting the role of histidine depletion-induced GCN2 activation regulating cholesterol biosynthesis, and the dependency on external cholesterol in T-ALL proliferation and survival.

## Cholesterol supplementation rescues the antileukemic effects of histidine restriction

Given the consistent finding that histidine restriction leads to downregulation in cholesterol biosynthesis at multiple levels, we next tested whether cholesterol supplementation might be able to rescue the therapeutic effects of histidine restriction. Interestingly, while cholesterol had no significant effect in the proliferation Jurkat cells grown in control RPMI media with 100% histidine levels, cholesterol supplementation resulted in a dose-dependent rescue of cell proliferation when cells were grown in media with reduced levels of histidine (**Fig. 6f**), demonstrating the importance of cholesterol in mediating the observed phenotype. Finally, we decided to test whether cholesterol supplementation *in vivo* would also rescue the strong antileukemic effects observed upon dietary histidine restriction in leukemic mice. Supplementation of histidine-restricted diets with 1.5% cholesterol, a level that has been previously used in multiple mouse studies^35,36^, led to a rescue of the decreased cholesterol serum levels induced by histidine restriction (**Fig. 6g**). More strikingly, while cholesterol addition to control diet did not impact leukemia progression, it led to a significant rescue of the antileukemic effects of histidine restriction, resulting in a significant acceleration of disease kinetics (**Fig. 6h**). Overall, these results unequivocally demonstrate that cholesterol downregulation partly mediates therapeutic effects of dietary histidine restriction in leukemia *in vivo*.

## Discussion

We performed comprehensive dietary amino acid dropout screens in primary NOTCH1-driven T-ALL to unbiasedly identify metabolic vulnerabilities. While dietary interventions show promise in preclinical models of cancer^11,12^ − including ketogenic diets^37–39^ or diets lacking certain amino acids, such as methionine^17,18^, valine^15^, serine/glycine^40^ or arginine/proline^13^ − most studies so far focus on the effects of single dietary interventions in immunodeficient mice. We strategically chose T-ALL given its clinical responsiveness to metabolic interventions^7^ and its allowing for high-throughput experiments in an orthotopic and immunocompetent setting. Our experiments confirmed the therapeutic effects of methionine or valine restriction but, most surprisingly, unveiled strong antileukemic effects for histidine restriction. Using diets with varying histidine levels, we observed dose-dependent oncogenic effects in leukemia progression *in vivo*, indicating that even diets with reduced (not eliminated) histidine levels may lead to therapeutic effects in patients. Histidine restriction therapeutic effects were confirmed in immunodeficient mice harboring human T-ALL PDXs with different driver mutations, indicating that this phenotype is at least partly tumor cell-intrinsic and reinforcing its potential for the treatment of human T-ALL.

Feeding healthy mice histidine-restricted diets for 2 weeks led to a dramatic impairment of normal T-cell development, in line with the effects of most therapeutic strategies in T-ALL. Prolonged feeding for over a month significantly decreased leukocyte populations in peripheral blood, with an expected progressive weight loss. Still, histidine restriction was fairly well tolerated in our experiments when compared to other diets lacking essential amino acids, such as valine- or leucine-free diets. These results imply an untapped potential for targeting histidine both in leukemia and in immunotherapy or immune-related diseases.

Our experiments to dissect the mechanistic effects of histidine restriction in T-ALL revealed an unexpected link between histidine deprivation and cholesterol biosynthesis suppression. Integrated transcriptomic, proteomic and ribosome profiling analysis showed histidine scarcity to cause ribosome stalling-mediated GCN2 activation to impair cholesterol biosynthesis across all expression levels (**Fig. 6i**). While the mechanism might be due to its impact on the master regulator SREBF2^41^, further mechanistic studies are needed to fully unravel this connection. The link between histidine availability and cholesterol metabolism represents a previously unrecognized regulatory axis. Critically, dietary cholesterol supplementation partially rescued the antileukemic effects of histidine restriction *in vivo*, mechanistically confirming that cholesterol pathway suppression mediates its therapeutic effects. Related to this, notably, rats fed with excess dietary histidine were shown to develop hypercholesterolemia in vintage studies^22,23^, overall underscoring a previously underrated relevance for the link between histidine and cholesterol. These results suggest that histidine-targeting strategies may be useful as novel anti-cholesterol therapies. They also reinforce the findings that cholesterol-targeting strategies show antitumor effects across multiple cancer types^42^, including a specific subtype of T-ALL^43^. This raises the possibility that treatment with cholesterol-lowering statins might have antileukemic effects in patients and/or might influence responses/relapses in adult T-ALL patients which might be concomitantly treated for hypercholesterolemia.

Our data suggests that restriction of most essential amino acids leads to the transcriptional downregulation of cholesterol biosynthesis. Thus, the stronger *in vivo* antileukemic effects with histidine restriction might be related to the specific organismal homeostasis of histidine, which makes its dietary depletion much better tolerated than that of leucine or valine. A relevant question is how broadly dietary histidine-restriction might be used in cancer treatment. While it is possible that histidine restriction shows antitumor effects across multiple cancer types, these will likely depend on the cell of origin and specific tumor type. Given its effects on normal leukocyte populations, it is tempting to speculate that histidine restriction will likely have therapeutic effects in other types of leukemia or lymphoma. Still, additional experiments encompassing other primary models of cancer (using both hematological malignancies but also solid tumors) are warranted to address this.

A previous study suggested a therapeutic role for histidine supplementation in combination with methotrexate treatment in an erythroleukemia cell line^44^. Erythroleukemia is a rare type of acute myeloid leukemia, leaving room for the possibility that histidine levels affect myeloid and lymphoid leukemias in opposing ways. Consistent with this hypothesis, the histidine catabolizing enzyme FTCD highlighted by that study is negligibly expressed in T-ALL (**Extended Data Fig. 11**), suggesting that this phenotype is specific for erythroleukemia. Additionally, Kanarek et al.^44^ addressed the effects of histidine only in the context of concomitant methotrexate treatment. Since histidine deprivation leads to cytostatic effects, and methotrexate primarily targets rapidly dividing cells^45,46^, histidine deprivation may impair cell cycle progression, negatively impacting on methotrexate effects.

For the clinical translation of our findings to T-ALL patients, a seemingly easy way to obtain therapeutic effects would be to carefully craft dietary recommendations that avoid histidine-rich food^7^. However, targeting a specific amino acid by dietary interventions alone is challenging. On the other hand, one of the core components of standard-of-care antileukemic regimens is the enzyme asparaginase, which catabolizes the amino acid asparagine^7^. Thus, we postulate that an analogous approach by an enzyme catabolizing histidine (e.g. histidine ammonia lyase)^47^, would likely lead to the strongest therapeutic effects. Experiments to specifically test this hypothesis and analyze the effects of histidine ammonia lyases from different origins, as well as the effects of free versus pegylated enzyme versions, are warranted.

Our comprehensive and unbiased experiments unveiled a previously unknown critical role for dietary histidine in leukemia progression through its impact on translational control and cholesterol metabolism, pioneering histidine-targeting strategies for therapeutic use in leukemia and other human diseases.

## Acknowledgements

We thank Adolfo A. Ferrando (Regeneron Genetics Center, New York) and Antonio Maraver (IRCM, Montpellier) for their constant constructive criticism and support, as well as Jan Müller (RNA Biochemistry laboratory, University Bern) for his technical guidance on ribosome profiling. We also thank everyone involved with JuanLord for their support. Mass spectrometry based proteomic analyses were performed by the Proteomics Research Infrastructure (PRI) at the University of Copenhagen (UCPH), supported by the Novo Nordisk Foundation (NNF) (grant agreement number NNF19SA0059305).

## Author contribution statement

K.M. performed most *in vivo* experiments and molecular biology assays. S.U. performed Ribo-seq, matched RNA-seq, proteomic profiling and most molecular biology assays. V.dS. performed multiple *in vivo* experiments, molecular biology experiments and provided guidance to K.M. C.T. and A.S. performed most computational analyses. O.K., M.A., G.C and T.K. performed some *in vitro* assays and helped with *in vivo* experiments. M.E. D-R. performed and analyzed all metabolomic experiments, supervised by X.S. E.P.W. and J.D.R. provided critical scientific input C.E. and S.L. co-supervised Ribo-Seq experiments. C.E. performed quality control of Ribo-seq. P.S. performed bioinformatic analysis and omics data integration and M.W. performed and analyzed proteomics. D.L. measured cholesterol levels. R.J.M. designed and supervised all Ribo-seq, proteomic profiling and omics data integration and provided overall scientific guidance. D.H. designed the study, supervised the research, attracted funding, provided scientific guidance and wrote the manuscript, together with input from all authors.

## Methods

### Pan-amino acid dietary restriction in T-ALL *in vivo*

Female C57BL/6 mice (Taconic Farms, Inc.), aged 8–10 weeks upon arrival, were acclimated to the local housing facility for 72 hours before experimentation. Amino acid-restricted diets were obtained from Research Diets, Inc., and were formulated based on a control diet (L-amino acid diet, A10021B). For each restriction, the specified amino acid was removed from the control formulation, with the remaining amino acids proportionally adjusted to maintain an equivalent total protein content.

### Mouse T-ALL survival studies

HDΔPEST-NOTCH1-GFP (1x10^6^) leukemia cells, previously established in primary mice, were transplanted into sub-lethally irradiated (4.5Gy) secondary recipients. 48 hours post-leukemic transplantation, recipient mice were subjected to either a control diet or individual amino-acid restricted diets, provided *ad libitum*.

For *in vivo* histidine dose-dependency, secondary recipients were subjected to a control diet or diets with decreasing amounts of histidine (50%, 25%, and 10% of control diet), with the remaining amino acids proportionally adjusted to maintain an equivalent total protein content. For leukemia acceleration studies, secondary recipients were subjected to a control diet or histidine-enriched diet containing a 10-fold standard histidine content while maintaining the total amino acid content constant.

For human PDX survival studies, PDTALL-10 or PDTALL-13 (1x10^6^) cells were injected into sub-lethally irradiated (2.5Gy) 8–10-week-old secondary NRG (JAX# 007799) recipients and fed with a control diet or a diet without histidine.

For the *in vivo* cholesterol rescue, secondary recipient mice were subjected to control diet or no histidine diet, either alone or supplemented with 1.5gm% dietary cholesterol, and survival analysis was performed.

All experimental mice were weighed twice weekly, monitored twice daily, and food intake was measured weekly by recording the amount of food provided at the beginning of the week and the amount remaining at the end of the week. Humane endpoints included signs of distress (e.g., hunched posture), significant weight loss, or loss of motor function.

### Hematological parameters

To evaluate the effect of histidine restriction on hematological parameters, 8-week-old healthy wild-type C57BL/6 mice were kept on control diet or no histidine diet for > 1 month. Their peripheral blood parameters were analyzed with a HESKA Element Ht5 analyzer.

### Cell culture and media preparation

Jurkat, DND41, HPB-ALL, MOLT3 and CUTLL1 cell lines were maintained in standard conditions in a humidified atmosphere at 5% CO_2_ at 37 °C in RPMI 1640 Media (HyClone, SH3002701, Fisher Scientific) supplemented with 10–20% FBS (Gemini Bio-Products, 900-108) and 1% penicillin–streptomycin (VWR, 45000-652).

Histidine restriction studies used histidine-free RPMI 1640 (USBio, R9000), prepared following the manufacturer’s instructions and reconstituted with L-histidine (H5100, USBio) at standard concentration (96.7 μM, control) or adjusted to 50% (48.35 μM), 25% (24.1 μM), 12.5% (12.08 μM), or 0% (no histidine).

For essential amino acid restriction experiments, branched-chain amino acid (BCAA)– free medium (USBio, R8999-20) was reconstituted with L-valine (Sigma, V0513), L-isoleucine (Sigma, I7403), and L-leucine (Sigma, L8912) at standard control (170.9 μM Valine, 381.6 μM Leucine, 381.6 μM Isoleucine) or 25% concentrations. Methionine-free medium (Fisher Scientific, A1451701) was supplemented with L-methionine (Sigma, M5308) at standard (100.6 μM, control) or 25% concentrations. Phenylalanine-free medium (USBio, R9000-01) was supplemented with L-phenylalanine (Sigma, P5482) at standard (90.9 μM, control) or 25% concentrations. Tryptophan-free medium (Teknova, R9940) was supplemented with L-tryptophan (Sigma, T8941) at standard (24.5 μM, control) or 25% concentrations.

### Custom media

For selected experiments, RPMI was prepared as a customized solution of all the single ingredients in-house. The formulation from manufacturer (Gibco, 11875085) was used as a template, and histidine was excluded from the base media for the purpose of metabolite dropouts. Histidine was added back using their 1000x concentrated solution either manually, or for the purpose of 384-well-plate screenings by Echo 655 liquid handler (Beckman-Coulter).

For cholesterol rescue experiments *in vitro*, lipid-depleted dialyzed FBS was used (10% supplementation). Lipid depletion was performed treating dialyzed FBS (Gibco, 26400044, lot 2675582P) a previously published method with fumed silica (Aerosil-380)^48^.

### Cell survival analyses

T-ALL cell lines were pre-treated for 24 h in RPMI supplemented with 10% dialyzed FBS and 1% penicillin–streptomycin, then plated in triplicate in 12-well plates (300,000 cells/well; 1 mL total volume) in media containing the indicated histidine concentrations. Cells were counted every 3 days using a Countess II FL instrument (Fisher Scientific).

In high-throughput settings, cells were seeded at a defined density (Jurkat 4000/well, DND41 and HPB-ALL at 8000/well) in a 384-well plate in 40 uL of media. At defined timepoints, cells were stained with CyQuant assay (Invitrogen, C7026) and imaged using high-throughput microscopy system Perkin-Elmer Operetta (method adapted from^49^). Images were analyzed in Perkin-Elmer Harmony software.

### Flow cytometry analyses

For T-cell development studies, healthy 6-8-week-old female C57BL/6 mice were placed on a control or no histidine diet for 2 weeks. Following euthanasia, thymi were collected and single-cell suspensions of total thymocytes were prepared by disrupting the thymus through a 70-mm filter. Red cells were removed by incubating in ammonium-chloride-potassium lysis buffer (155mM NH4Cl, 12mM KHCO3 and 0.1mM EDTA) for 5 min at room temperature. Single cells were stained with: CD4 PE-eFluor 610 (1:400, RM4-5, Thermo Fisher), CD8a PE (1:200, 53-6.7, BD Pharmingen), CD44 PerCP–Cy5.5 (1:400, IM7, Thermo Fisher), and CD25 Alexa Fluor 488 (1:1000, 7D4, Thermo Fisher). Thymocyte populations were gated as CD4 SP (CD4⁺CD8a⁻), CD8 SP (CD4⁻CD8a⁺), DP (CD4⁺CD8a⁺), and DN (CD4⁻CD8a⁻). DN cells were further sub-categorized into DN1 (CD44⁺CD25⁻), DN2 (CD44⁺CD25⁺), DN3 (CD44⁻CD25⁺), and DN4 (CD44⁻CD25⁻) stages.

Cell cycle and apoptosis in Jurkat cells were analyzed at day 3 using PI/RNase staining buffer (BD Pharmingen, 550825), and apoptosis quantified using PE–Annexin V Apoptosis Detection Kit I (BD Pharmingen, 559763). Data was acquired on an Attune NxT flow cytometer (Thermo Fisher) and analyzed in FlowJo v10.6.2 (BD).

### Analysis of active protein synthesis by puromycin incorporation

Jurkat cells cultured in control or 25% histidine media for 72 h were pulsed with puromycin (5 μg/mL) for 15 min at 37 °C, stained with LIVE/DEAD™ Fixable Violet Dead Cell Stain (Invitrogen, L34963), fixed (BD Cytofix/Cytoperm™, 554714), stained intracellularly with anti-puromycin (BioLegend, clone 2A4, mouse, 381504), and analyzed by flow cytometry. Alternatively, for western blotting cells were cultured in custom Histidine-free RPMI supplemented with 10% dialyzed FBS and different levels of Histidine (100%, 50%, 25% and 12.5%). After 72 hours, cells were exposed to puromycin (5 µg/mL) for 15 minutes (SUNSET protocol)^50^. Then cycloheximide was added to final concentration 100µg/mL to inhibit translation. Lysates were separated electrophoretically and transferred to PVDF membrane (Bio-Rad), blocked with milk buffer, and detected with a mouse-monoclonal anti-Puromycin antibody (clone 12D10, #MABE343, Milipore) at 1:1000 at 4°C overnight followed by goat anti-mouse secondary antibody at a 1:20,000 dilution.

### Western blotting

Cellular proteins were extracted from Jurkat cells and DC Protein Assay Kit (Bio-Rad) was used to quantify total protein. Homogenates containing equal amount of protein were denatured with 4 × loading buffer, separated on a 4-20% polyacrylamide gel and electroblotted onto PVDF membranes. The membranes were blocked with 5% skimmed dried milk in TBS-T, or for phospho-proteins with 5% BSA in TBS-T, at room temperature for 1.5 h, incubated with primary antibodies overnight at 4 °C. The following primary antibodies were used:

| Target | Dilution | Source | Product no. | Lot No. |
| --- | --- | --- | --- | --- |
| ATF4 | 1:800 | Proteintech | 10835-1-AP | 00146594 |
| PERK | 1:4000 | Proteintech | 20582-1-AP | 00101749 |
| p-PERK | 1:15000 | Proteintech | 82534-1-RR | 23006222 |
| XBP1-s | 1:1000 | Cell Signalling Technology | 40435 | 5 |
| ATF6 | 1:1000 | Cell Signalling Technology | 65880 | 5 |
| HMGCS1 | 1:1000 | Proteintech | 17643-1-AP | 00024906 |
| HMGCR | 1:1000 | abcam | CL0260 | 1075659-14 |
| SREBF2 | 1:4000 | Proteintech | 28212-1-AP | 00115966 |
| GCN2 | 1:600 | Proteintech | 27944-1-AP | 00166737 |
| p-GCN2 | 1:1000 | Cell Signalling Technology | 94668 | 4 |
| Anti-Puromycin clone 12D10 | 1:1000 | Sigma-Aldrich | MABE343 | 4191133 |
| p-mTOR | 1:1000 | Cell Signalling Technology | 5536 | 12 |
| p-S6K | 1:1000 | Cell Signalling Technology | 9234 | 16 |
| p-4EBP1 | 1:1000 | Cell Signalling Technology | 2855 | 30 |
| 4EBP1 | 1:1000 | Cell Signalling Technology | 9644S | 12 |
| actin | 1:20000 | Proteintech | HRP-66009 | 21009669 |
| GAPDH | 1:20000 | Proteintech | HRP-60004 | 21010937 |

Secondary antibodies coupled to HRP (Proteintech: anti-Rabbit #RGAR001, lot 2000851; anti-Mouse #RGAM001, lot 20000844) were used to detect antibody binding by chemiluminescence. Antigen-antibody signals were visualized using the Clarity Wester ECL Solution (Bio-Rad, 170-5061), according to the manufacturer’s protocol using the Bio-Rad imaging system.

### U-^13^C-histidine labeling

Jurkat cells were pre-treated for 24 h in RPMI with 10% dialyzed FBS and 1% penicillin– streptomycin, then plated at 5 × 10⁶ cells/well (3 mL) in histidine-free RPMI containing 10% dialyzed FBS, 1% penicillin–streptomycin, and U-^13^C–L-histidine (96.7 μM, 100%; Cambridge Isotope Laboratories). Labeling was performed for 4 h or 10 days.

### Metabolite extraction and metabolomic analyses

Cell suspensions were centrifuged (400 × g, 2 min, RT), washed with PBS, and extracted with 1 mL ice-cold methanol:acetonitrile:water (40:40:20, v/v/v) + 0.5% formic acid. After 5 min on ice, extracts were neutralized with 50 μL 15% ammonium bicarbonate and centrifuged (15,000 × g, 10 min, 4 °C). Supernatants were snap-frozen for LC-MS. Media metabolomics used 20 μL culture medium diluted in 980 μL extraction solvent, neutralized with 80 μL ammonium bicarbonate.

### RNA-seq analyses

Total RNA was isolated using the QIAshredder (Qiagen, 79656) and RNeasy Mini Kit (Qiagen, 74106), following the manufacturer’s protocols. RNA library preparation and next-generation sequencing were performed on the Illumina NextSeq platform.

### High-throughput nutrient dependency and drug response profiling

Using a high-throughput liquid handling system ECHO-655 (Beckman Coulter), the amino acids were added in a concentration gradient manner into 384-well plate (Greiner, 781091) prefilled with 20 uL media. Cells were collected by centrifugation, resuspended in customized RPMI media (lacking Histidine) supplemented with 10% dialyzed FBS (or lipid-depleted dialyzed FBS for Cholesterol rescue). 20 uL of cell suspension were added per well. When indicated, another compound (GCN2iB (MedChemExpress, HY-112654), NXP800 (MedChemExpress, HY-145927) or cholesterol water-soluble (Sigma, C4951)) was added into a matrix. Cells were cultured in the plates at 37°C 5% CO2 for the indicated time. For imaging, cells were stained with CyQuant solution (Invitrogen, C35011), incubated for 1 hour and imaged with Perkin-Elmer Operetta with 10x objective. Using the Harmony 4.9 software the images were analyzed to recognize and count only the live lymphoblasts for further analyses.

### Ribosome profiling

Ribosome profiling was performed as described previously^51,52^. Jurkat cells were seeded in 50 mL of customized RPMI with 10% dialyzed FBS and 100% (96.77 µM) or 25% (24.19 µM) histidine at a density of 3 x 10^5^ cells/mL. Aliquots for cell counts were harvested, and cells were counted using Trypan blue. Cells were collected by centrifugation (5 min at 300g), and the pellets were snap-frozen in liquid nitrogen.

Ice-cold Human Lysis Buffer (10 mM Tris-HCl, pH 7.5, 100 mM NaCl, 5 mM MgCl2, 1% Triton X-100, freshly added 0.5% sodium deoxycholate, 1 mM DTT, 100 µg/mL cycloheximide, as previously described^51^) was added to the cell pellet (500 µL per 90 million cells). Lysates were resuspended, transferred to Eppendorf tube and centrifuged at 10,000x g for 5 min at 4°C. The supernatant was collected in a clean tube and snap-frozen in liquid nitrogen.

Samples were thawed and RNA concentration was measured using Nanodrop system. Equal amounts of 300 ug total RNA for each sample was collected, subjected to digestion with 12 U of Turbo DNase (Invitrogen™ AM2239) and 1 000 U RNase I (Invitrogen™ AM2295). Polysomes were digested for 1 hour at 22°C at 1,400 rpm, followed by 200 U SUPERase·In™ (Invitrogen™ AM2694) and sodium deoxycholate to final concentration of 0.5%. Monosomes were recovered by ultracentrifugation at 35,000 rpm in a cooled SW41 Ti rotor over a sucrose gradient (10 – 50 % sucrose in polysome buffer containing 100 ug/mL CHX and 1 mM DTT). RNA was extracted using heated P/C/I (phenol/chloroform/isoamylalcohol) solution. The RNA was subjected to electrophoresis under denaturing conditions in 15% PAA/1xTBE/8M-Urea gel. Ribosome protected fragments (RPFs) of 28-32 nucleotides were eluted from the gel in RNA-extraction buffer (0.3 M NaOAc, 1 mM EDTA pH 8.0, 0.25% SDS) at 4°C overnight. After filtration, precipitation was performed using ∼300 uL eluent, 4 uL glycogen (5 mg/mL) and 900 µL EtOH. RPFs were dephosphorylated using T4 PNK (NEB) enzyme, 10 U per reaction. Subsequently, fragments were ligated to 0.5 µg adenylated cloning linker using 200 U of T4 RNA ligase 2, truncated KQ (NEB). Subsequently, reverse transcription (RT) was performed with 2,000 U of SuperScript III (Invitrogen™) followed by a size-selection of the RT product on 10% PAA/1x TBE/8M-Urea gel. The RT product was depleted of ribosomal RNA contaminants using a set of biotinylated oligonucleotides. For rRNA depletion, cDNA was mixed with biotinylated subtraction oligo pool (10 µM each) and 20× SSC, denatured at 100 °C for 90 s, and annealed by cooling at 0.1 °C/s to 37 °C, followed by incubation for 15 min at 37°C. The reaction was then combined with 25 µL of pre-washed MyOne Streptavidin C1 Dynabeads (Invitrogen, 65001) and incubated at 37°C for 15 min (1,000 rpm). Beads were magnetically separated, and the supernatant was recovered and ethanol-precipitated. Next, samples were subjected to circular ligation using 100 U of CircLigase (LGC Biosearch Technologies, CL9025K) and amplified by PCR using Phusion polymerase (NEB) for adjusted number of cycles of denaturation (15 s at 98°C), annealing (30 s of 62°C) and elongation (15 s at 72°C) using primer pairs compatible with the Illumina sequencing platform. The resulting ribosome-profiling libraries were sequenced on a NextSeq 1000 instrument (Illumina).

Total RNA was isolated from the same lysates used to obtain RPFs. 1 mL of TRIzol Reagent (Invitrogen, 15596026) reagent was added to 55 µL of the lysate (containing material from ∼10 M cells), mixed by vortexing and incubated for 5 minutes at 4°C. Then, 0.2 volumes of BCP were added and the content was shaken for 15 s and incubated for 3 min at RT. Samples were spun at 10,000xg for 10 min at 4°C. Aqueous phase was collected, 1 mL of isopropanol was added, and the mixture was incubated at -20°C for 1 hour. Then the samples were centrifuged at 10,000xg for 10 min at 4°C. Pellets were washed with 75% EtOH and re-centrifuged briefly. The pellet was air-dried for 5 minutes and resuspended in 20 µL Nuclease-free H2O (Promega, P119E). The yield ranged between 1,000 and 1,6000 ng/µL.

### Proteomics

For total proteome sample preparation, cells were seeded in the customized media containing 100% or 25% Histidine (relative to RPMI content) at a density of 3x10^5^/mL. After 72 hours, 4 replicates per condition were harvested by centrifugation, washed by PBS and the pellet was snap frozen in liquid nitrogen and shipped on dry ice. Cell pellets were lysed in 120 µL Thermo Lysis Buffer supplemented with 2 µL Universal Nuclease. Lysates were homogenized by pipetting and heated at 95°C for 10 min, followed by centrifugation to remove insoluble material. Protein concentrations were determined using the Pierce BCA assay. Proteins were reduced with 5 mM TCEP for 15 min at 55 °C and alkylated with 20 mM CAA for 30 min at room temperature. Digestion was performed overnight at 37 °C using trypsin and LysC at a 1:50 enzyme-to-protein ratio. Resulting peptides were acidified with 10% formic acid (FA) and desalted using a Thermo desalting plate. Peptides were eluted with 150 µL 40% acetonitrile (ACN) containing 5% ammonium hydroxide and dried in a SpeedVac concentrator at 45 °C.

For LC-MS Measurements, peptides were separated on a 50 cm μPAC NEO HPLC column (Thermo Scientific) operated at 50°C. Solvent A was 0.1% formic acid (FA) in water, and solvent B was 80% acetonitrile (ACN) with 0.1% FA. Separation was performed using a two-step gradient: 4–22.5% B over 40 min at 0.35 µL/min, followed by 22.5–45% B over 13 min at 0.35 µL/min. The total run time was 60 min including wash. Eluted peptides were introduced via an EASY-Spray source with a 30 µm steel emitter (Evosep) into a Tribrid Ascend mass spectrometer (Thermo Scientific) with a spray voltage of 2200 V.

Data were acquired in data-independent acquisition (DIA) mode. Full MS scans were recorded at 60,000 resolution across an m/z range of 400–900, with an AGC target of 250% and maximum injection time set to auto. DIA windows consisted of 42 scans with 12 Th isolation width and 1 Th overlap, covering the 400–900 m/z range. MS/MS scans were acquired at 15,000 resolution with an AGC target of 1000% and maximum injection time of 27 ms. Fragmentation was performed by higher-energy collisional dissociation (HCD) with normalized collision energy (NCE) of 30%.

### Data analysis

#### Reference sequences and annotations

The reference human genome (GRCh38; annotation version 113) and transcriptome were obtained from ENSEMBL^53^. For ribosome profiling, coding sequences (CDS) were curated in BioMart with the filters (a) Source (transcript): ensembl_havana (b) Transcript type: lncRNA, protein_coding (c) Ensembl Canonical: Only; and attributes (a) Transcript stable ID (b) Coding sequence (c) Upstream flank [18] (d) Downstream flank [18]. Sequences for rRNA and tRNA were obtained from NCBI (Nucleotide) and Genomic tRNA database, respectively^54,55^. Human proteome (reviewed and canonical) was obtained from UniProt^56^. Gene sets (*c2.cp.reactome.v2025.1.Hs.symbols.gmt*) for GSEA were obtained from MSigDB^57^.

#### Bulk RNA sequencing

Decoy (full genome) aware indexing of transcriptome was done using Salmon with the parameter “*--kmerLen 31*”^58^. The paired-end reads from bulk RNA-seq datasets were processed to remove the Illumina universal adapter (AGATCGGAAGAGCACACGTCTGAACTCCAGTCA) using Cutadapt (v4.4) with the parameters *“--poly-a -q 25 -m 25 -a AGATCGGAAGAGCACACGTCTGAACTCCAGTCA -A AGATCGGAAGAGCGTCGTGTAGGGAAAGAGTGT*”^59^. The resulting filtered reads were quantified against the index using Salmon with the parameters “*--libType A –gcBias --seqBias –useVBOpt --validateMappings*”^58^. Subsequent data analysis was performed with custom scripts in R. Data wrangling and visualization was performed using tidyverse package (v2.0.0)^60^. Transcript quantifications were imported using Tximeta and differential expression analysis was conducted using DESeq2 (v1.44)^61,62^. Cluster of genes changing expression with histidine concentrations were identified by likelihood ratio test using the function *DESeq(test = “LRT”, reduced = ∼ 1)*, where full model was ∼ condition (i.e. samples belonging to different histidine concentration). Differentially expressed genes (DEGs) were defined as having adjusted p-value < 1e-06. Genes in upregulated and downregulated clusters were further analyzed for functional enrichment using gProfiler2 (v0.2.3) (Bioconductor.org)^63^.

#### Ribosome profiling

Single-end reads were processed to remove adapter (*--quality-cutoff 30 --minimum-length 25 --cut 3 --no-indels --discard-untrimmed --adapter CTGTAGGCACCATCAAT*) and randomized nucleotides (*--cut -6 --quality-cutoff 30 --minimum-length 25*) using Cutadapt (v4.4)^59^. Resulting filtered reads were mapped to rRNA and tRNA using bowtie (v1.3.1) with the parameters “*--best --sam --un*”^64^. Reads that failed to align rRNA and tRNA were mapped to coding sequence using bowtie (v1.3.1) with the parameters “*—best --sam -m 1 -v 1 --norc --strata*”^64^. Coordinate-sorted alignment files were indexed using SAMtools (v1.17)^65^. Differential expression analysis was conducted using DESeq2 (v1.44)^61^. Differentially expressed genes (DEGs) were defined as having absolute log fold change > 1 and adjusted p-value < 0.05 (Wald test).

Codon occupancy was calculated using *ribomala* package (v0.5.4)^66^. Briefly, read length of interest (generally 28-31 nt) were determined based 3-nt periodicity on the CDS. For each read length, E-, P- and A-site were determined based on the highest ribosome footprint signal upstream of the start site. For each coding sequence, 20 codons from the start and end were excluded, and read counts (e.g. A-site) for each codon were divided by mean across the coding sequence to get relative enrichment. Coding sequences with mean read count of < 0.1 reads/codon were also excluded from the analysis.

#### Proteomics

Raw mass spectrometry data was processed using Spectronaut (v20.0) using default parameters^67^. The output was further analyzed using MSstats package (v4.16.1)^68^. Differentially expressed genes proteins were defined as having absolute log fold change > 0.585 and adjusted p-value < 0.05.

#### GSEA

GSEA was performed using clusterProfiler (v4.16.0) and enrichplot (v1.28.4)^69,70^. Genes and proteins were ranked based on shrunken LFC (RNA-seq and ribosome profiling) and LFC (proteomics), respectively^71^.

#### Data and code availability

The RNA-Seq and Ribo-Seq data are accessible under the accession number GEO: GSE313291. RNA-seq for Jurkat, DND41 and all EAA restriction experiments are accessible under GSE300867, GSE301665, GSE301669. The mass spectrometry proteomics data will be deposited to the ProteomeXchange Consortium via the PRIDE partner repository with the dataset identifier: PXD071870. No customized code has been developed for analysis. Additional information required to reanalyze the data reported in this manuscript is available from the corresponding author upon request.

