## Supplemental Figures for "A dietary pan-amino acid dropout screen *in vivo* reveals a critical role for histidine in T-ALL"

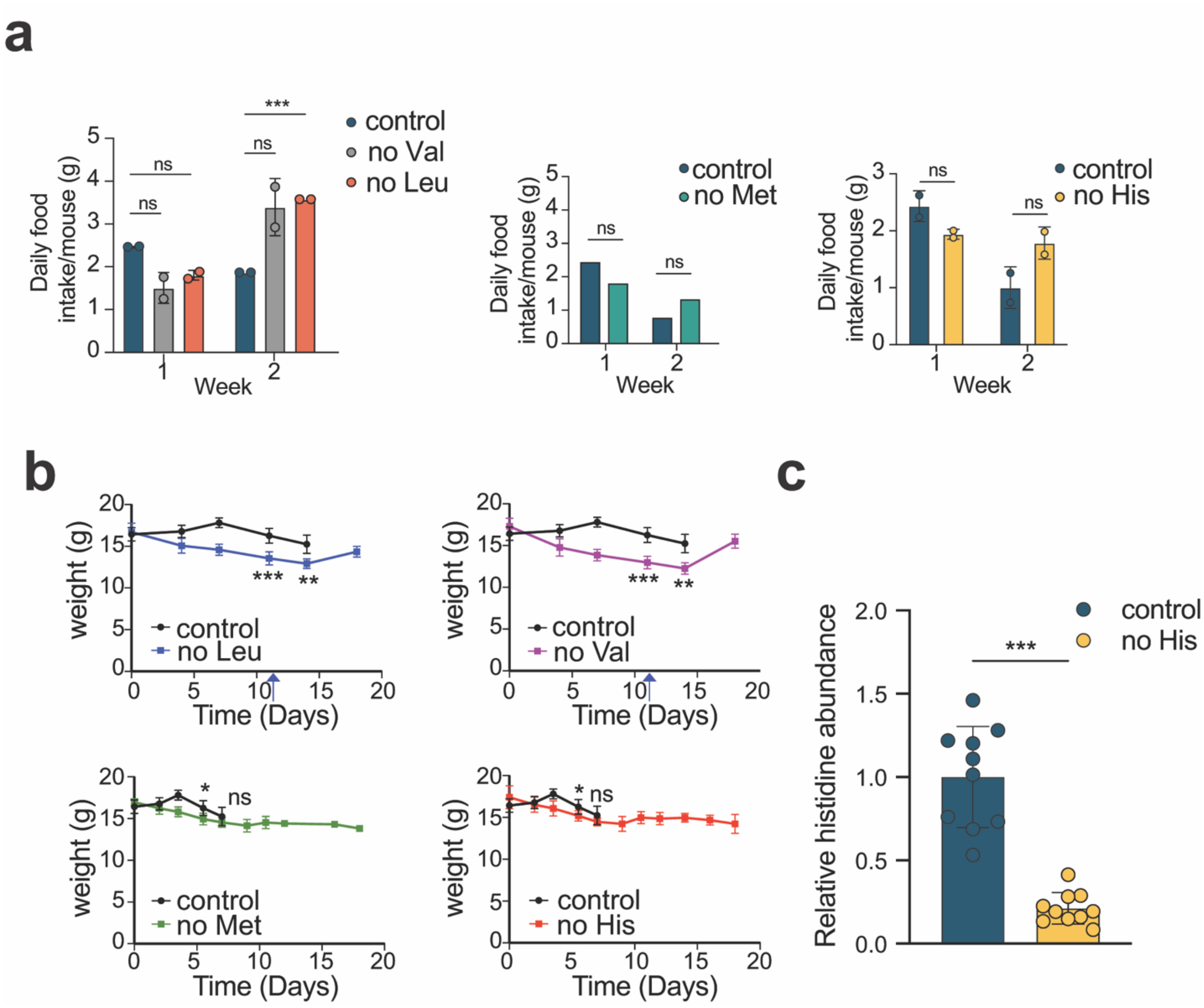


**Extended Data Fig. 1 │ Diets lacking histidine and methionine are well tolerated in leukemic mice. a**, Average diet consumption in leukemic mice during survival analyses. Diet intake was calculated weekly by measuring the amount of food provided at the beginning of the week and the remaining amount at the end of the week. **b**, Body weight of mice harboring NOTCH1-induced leukemias and fed with diets lacking leucine, valine, methionine or histidine. Arrow depicts switch to control diet due to acute weight loss. Data are mean ± sem (n=6 per group). **c**, Levels of histidine in the serum of leukemic mice after a 2-week feeding with normal chow or diet without histidine. *P* values in a–c were calculated using student's t-test; **P* < 0.05; ***P* < 0.01; ****P* < 0.005 (n=10 per group).

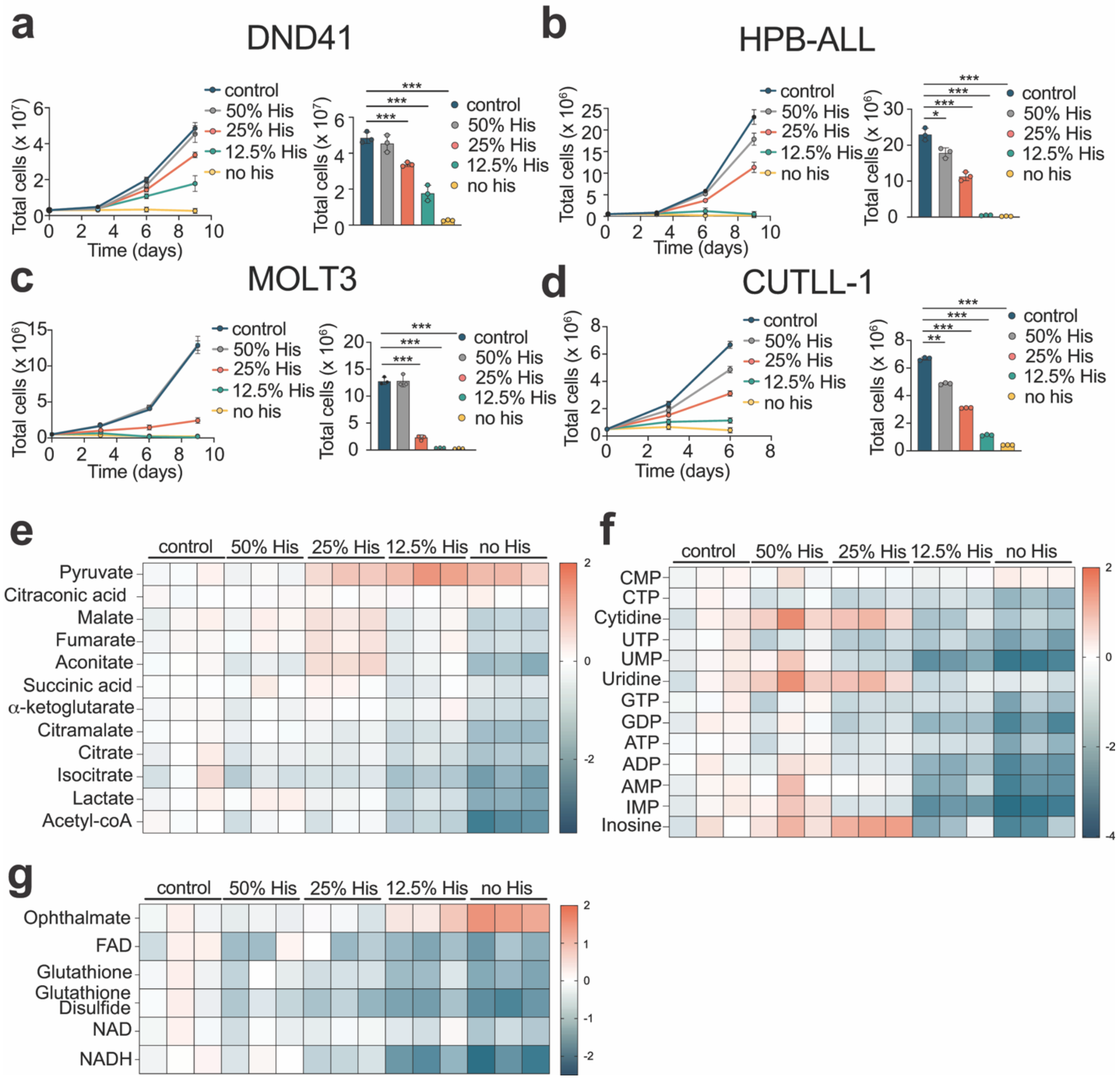


**Extended Data Fig. 2 | Histidine restriction shows antiproliferative effects and leads to significant metabolic changes *in vitro*. a–d**, Proliferation curve (left) and quantification at day 6 or 9 (right) of DND41 (**a**), HPB-ALL (**b**), MOLT3 (**c**) and CUTLL-1 (**d**) human T-ALL cells grown in normal RPMI media, RPMI with 50%, 25% or 12.5% levels of histidine, or RPMI without histidine. Data are mean ± sem (n=3). **e–g**, Heatmap showing differential intracellular abundances ($\log_2$) of TCA cycle intermediates (**e**), nucleotide intermediates (**f**) or redox metabolism intermediates (**g**) in Jurkat cells grown in normal RPMI media, RPMI with 50%, 25% or 12.5% levels of histidine, or RPMI without histidine for 3 days. *P* values in a–d were calculated using one-way ANOVA for multiple comparisons; *$P < 0.05$; **$P < 0.01$; ***$P < 0.005$.

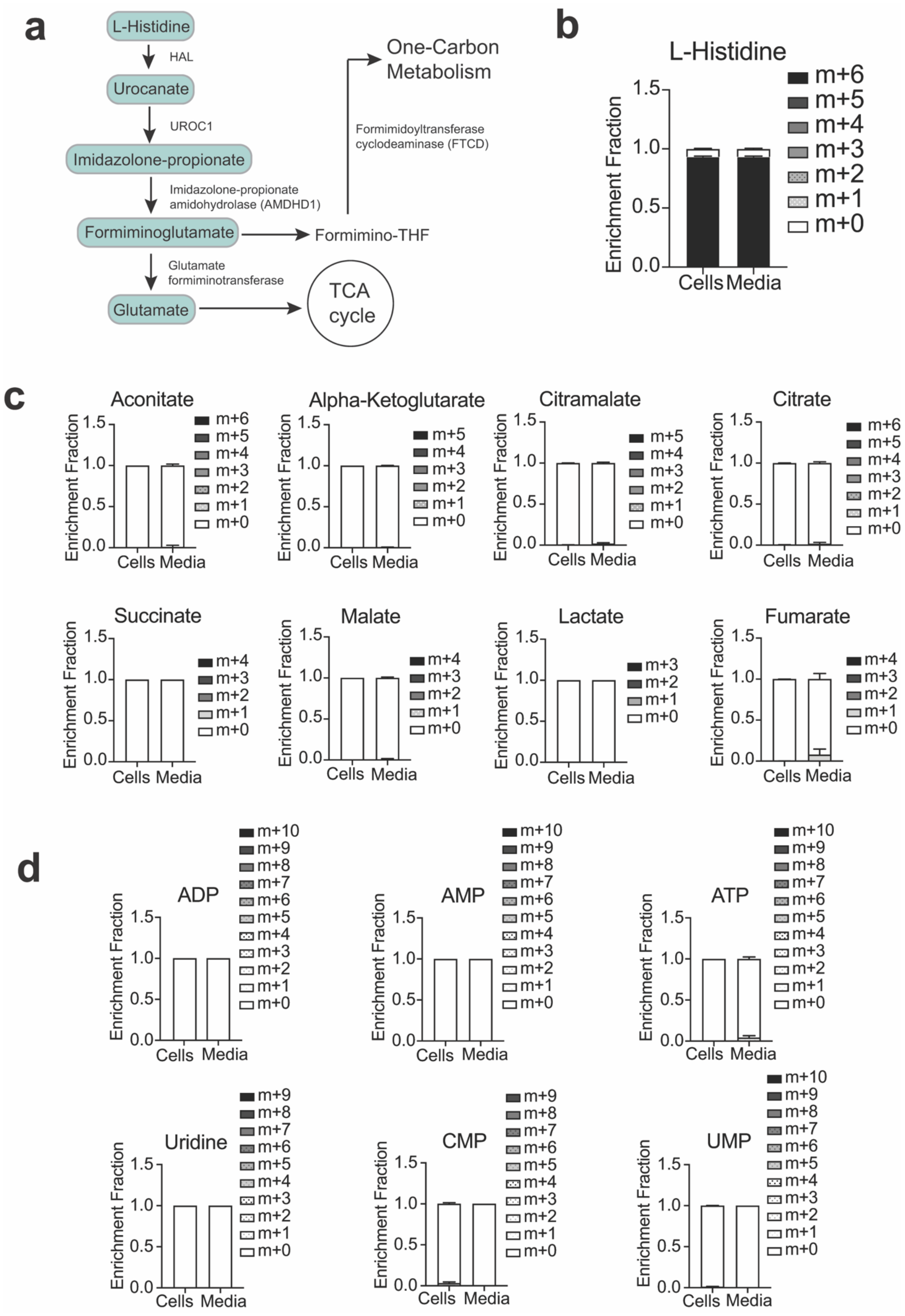
a
L-Histidine
HAL
Urocanate
UROC1
Imidazolone-propionate
Imidazolone-propionate amidohydrolase (AMDHD1)
Formiminoglutamate
Formimino-THF
Formimidoyltransferase cyclodeaminase (FTCD)
One-Carbon Metabolism
Glutamate formiminotransferase
Glutamate
TCA cycle
b
L-Histidine
Enrichment Fraction
m+6 m+5 m+4 m+3 m+2 m+1 m+0
Cells Media
c
Aconitate
Alpha-Ketoglutarate
Citramalate
Citrate
Succinate
Malate
Lactate
Fumarate
Enrichment Fraction
Cells Media
d
ADP
AMP
ATP
Uridine
CMP
UMP
Enrichment Fraction
m+10 m+9 m+8 m+7 m+6 m+5 m+4 m+3 m+2 m+1 m+0
Cells Media

**Extended Data Fig. 3 | Short-term *in vitro* stable isotope tracing of U-$^{13}$C-Histidine in Jurkat cells. a**, Schematic of the histidine degradation pathway. **b**, $^{13}$C-labeling pattern of histidine after 4 hours of U-$^{13}$C-histidine labeling in Jurkat cells in triplicates. **c**, $^{13}$C-labeling patterns of indicated metabolites after 4 hours of $^{13}$C-histidine labeling in Jurkat cells in triplicates. Data are mean ± sem (n=3).

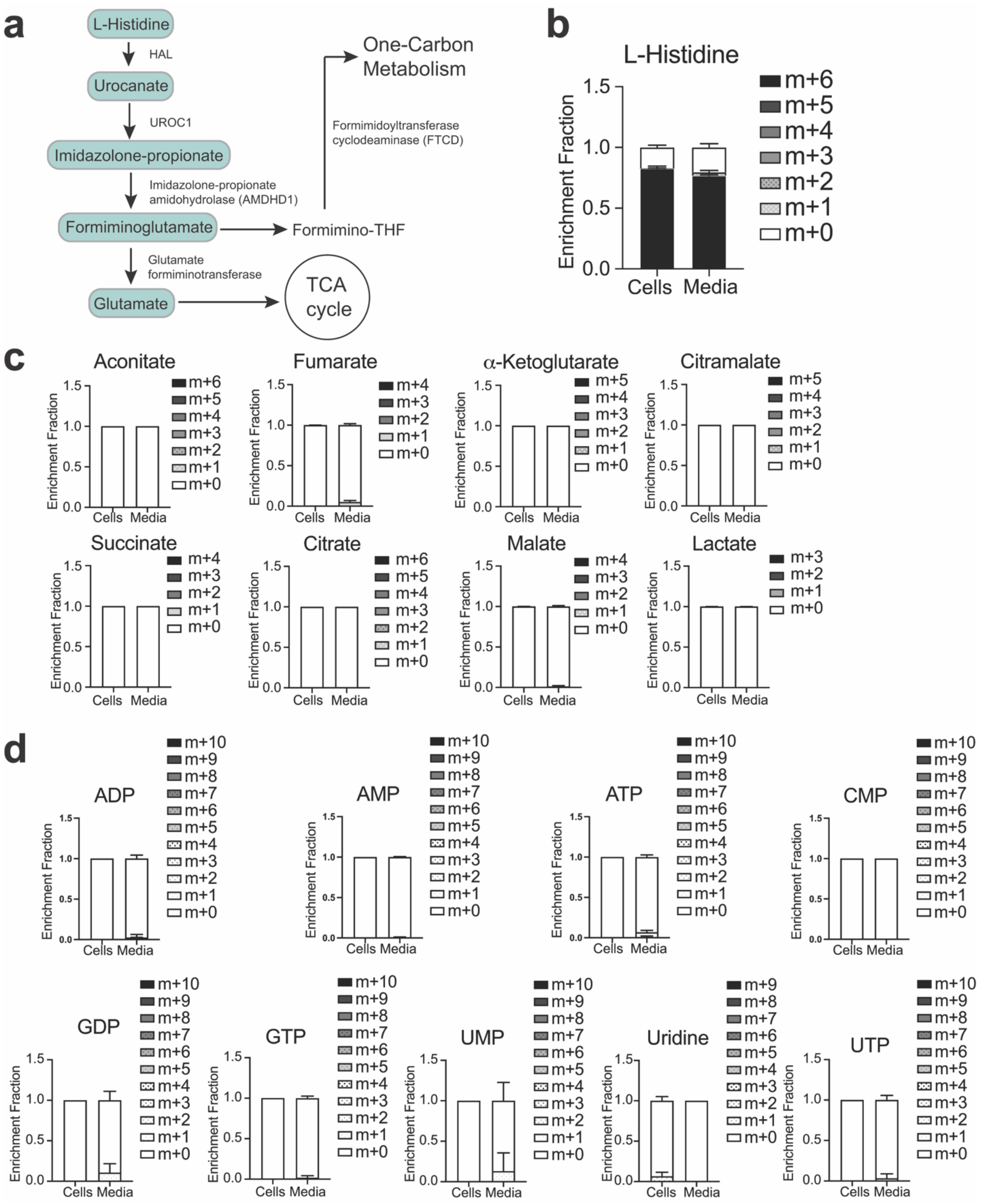


**Extended Data Fig. 4 │ Long-term *in vitro* stable isotope tracing of U-$^{13}$C-Histidine in Jurkat cells. a**, Schematic of histidine degradation pathway. **b**, $^{13}$C-labeling pattern of histidine after 10 days of U-$^{13}$C-Histidine labeling in Jurkat cells in triplicates. **c**, $^{13}$C-labeling patterns of indicated metabolites after 10 days of $^{13}$C-Histidine labeling in Jurkat cells in triplicates. Data are mean ± sem (n=3).

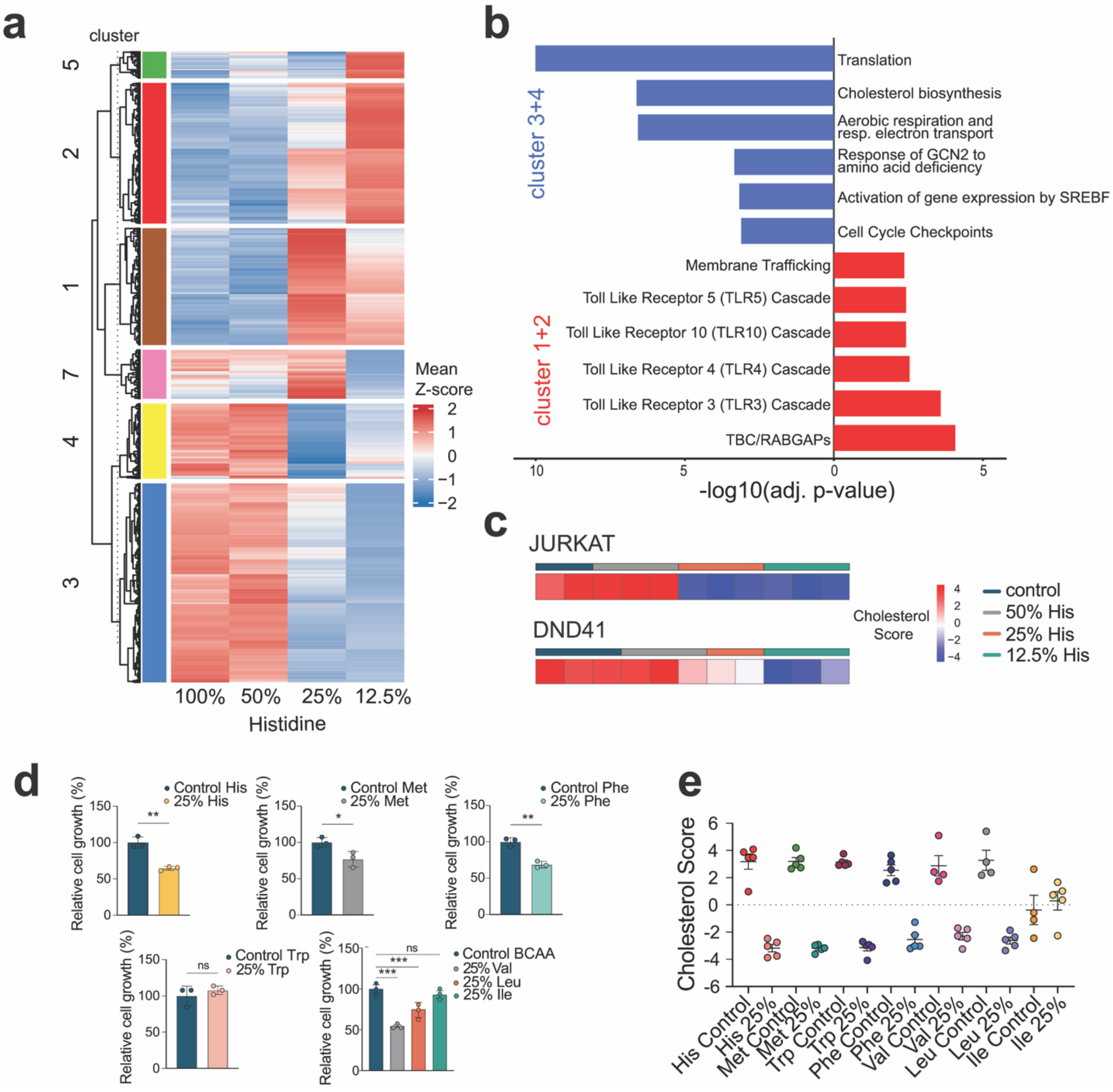


**Extended Data Fig. 5 | Transcriptional responses to histidine depletion in human T-ALL cells**. **a**, Hierarchical clustering of differentially expressed genes across histidine concentrations (100%, 50%, 25%, 12.5%) in DND41 cells cultured for 6 days (144 hours). Seven distinct clusters were identified based on gene expression Z-scores, revealing coordinated transcriptional programs responsive to histidine availability. **b**, Pathway analysis using gProfiler2 with the list of significantly down/upregulated genes (g:SCS method; P<0.01, log2FC >0.584). Clusters 3 and 4 (blue) are enriched for, cholesterol biosynthesis, GCN2 response and cell cycle–related pathways, whereas clusters 1 and 2 (red) show enrichment for membrane trafficking and toll like receptor signaling. **c**, Comparison of cholesterol biosynthesis pathway activity between Jurkat and DND41 cells using cholesterol pathway scores (n=3 per group). **d**–**e**, Proliferation (**d**) and cholesterol score (**e**) obtained upon culture of Jurkat cells for 72h in normal RPMI, or RPMI with 25% the normal levels of His, Met, Trp, Phe, Val, Leu or Ile (n=4-5 per group; data are mean ± sem).

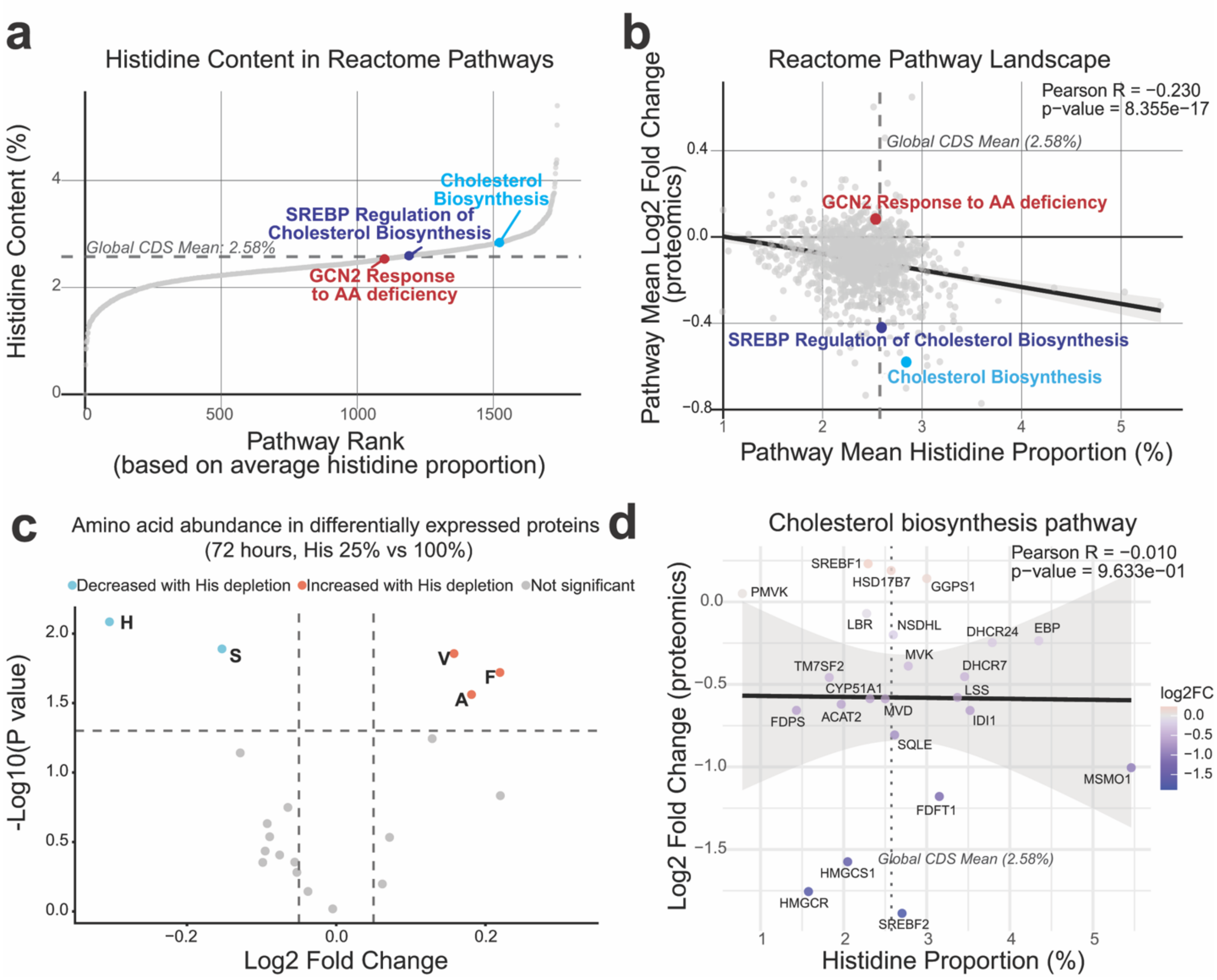


**Extended Data Fig. 6 | Histidine content has low impact on protein expression under histidine depletion. a**, Global ranking of Reactome pathways by histidine content; highlighted pathways of interest show no outlier distribution. **b**, Weak negative correlation (Pearson $R$ = –0.23) between pathway-mean histidine content and proteomic changes after 72 h (adapted from[1]). **c**, Downregulated proteins under histidine depletion are enriched for histidine codons (adapted from[2]). **d**, Lack of correlation (p-value > 0.05) between histidine content and expression for individual enzymes in the Cholesterol Biosynthesis pathway, indicating complex regulation.

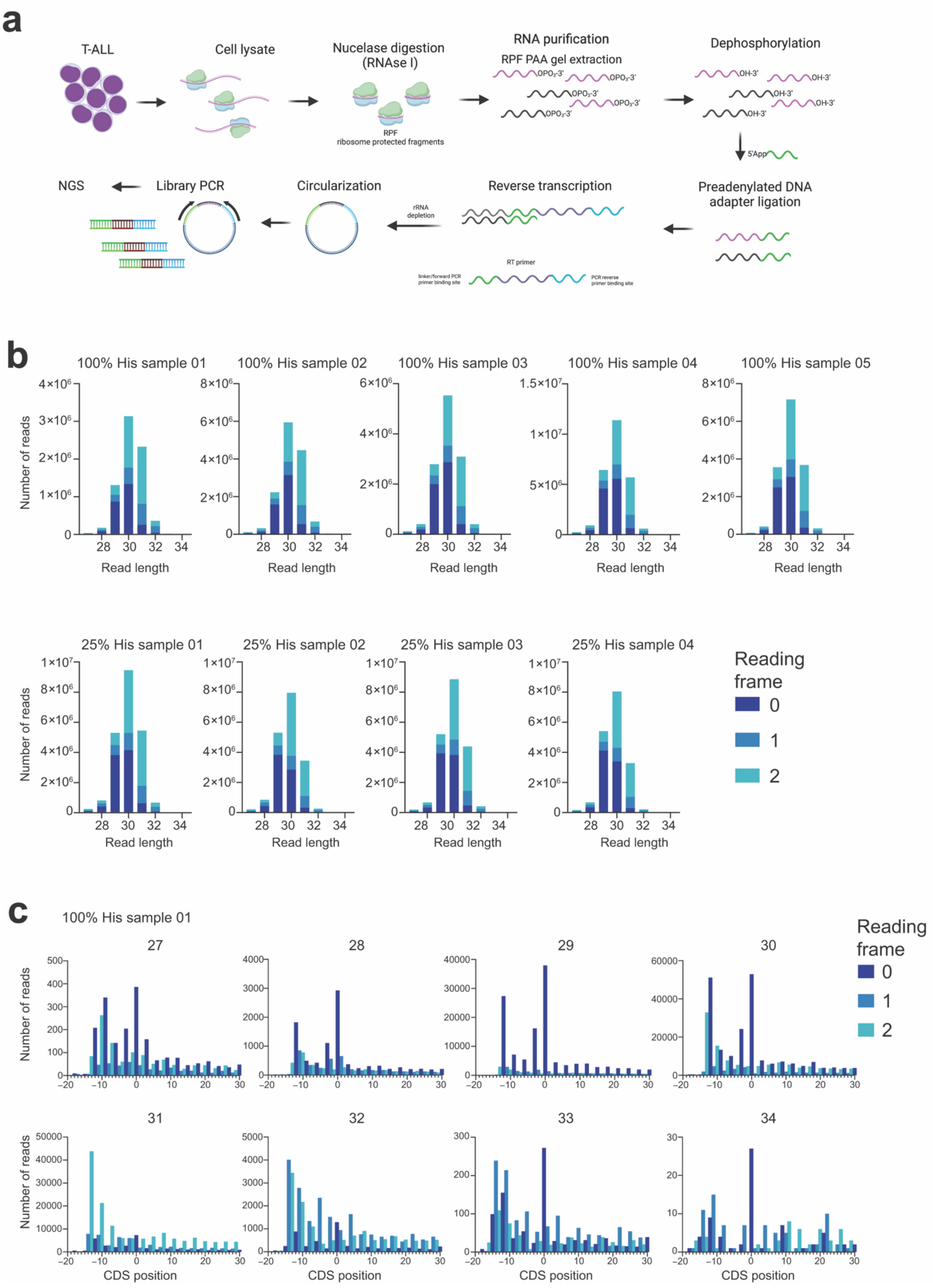
a
T-ALL
Cell lysate
Nucleation digestion (RNAse I)
RPF
ribosome protected fragments
RNA purification
RPF PAA gel extraction
Dephosphorylation
5'App
Preadenylated DNA adapter ligation
Reverse transcription
RT primer
linker/forward PCR primer binding site
PCR reverse primer binding site
rRNA depletion
Circularization
Library PCR
NGS
b
100% His sample 01
100% His sample 02
100% His sample 03
100% His sample 04
100% His sample 05
25% His sample 01
25% His sample 02
25% His sample 03
25% His sample 04
Number of reads
Read length
Reading frame
0
1
2
c
100% His sample 01
27
28
29
30
31
32
33
34
Number of reads
CDS position
Reading frame
0
1
2

**Extended Data Fig. 7 | Quality control of ribosome profiling data. a**, Schematic workflow of the ribosome profiling experiment (adapted from[3]) performed for Jurkat cells cultured in histidine 100% and histidine 25% media for 72 hours. Cell lysates were digested with RNase I to generate ribosome-protected fragments (RPFs), which were purified, dephosphorylated, ligated to pre-adenylated adapters, reverse-transcribed, circularized, and amplified for next-generation sequencing. **b**, Distribution of RPF read lengths across samples cultured in histidine-replete (100% His) or histidine-depleted (25% His) media. The characteristic enrichment of 28–30 nt fragments and the strong in-frame periodicity (reading frame 0) confirm high library quality and successful capture of translating ribosomes. **c**, Metagene analysis of read frame periodicity for RPFs of different lengths in a representative sample (100% His). Triplet periodicity centered on the start codon demonstrates correct P-site assignment and expected codon phasing.

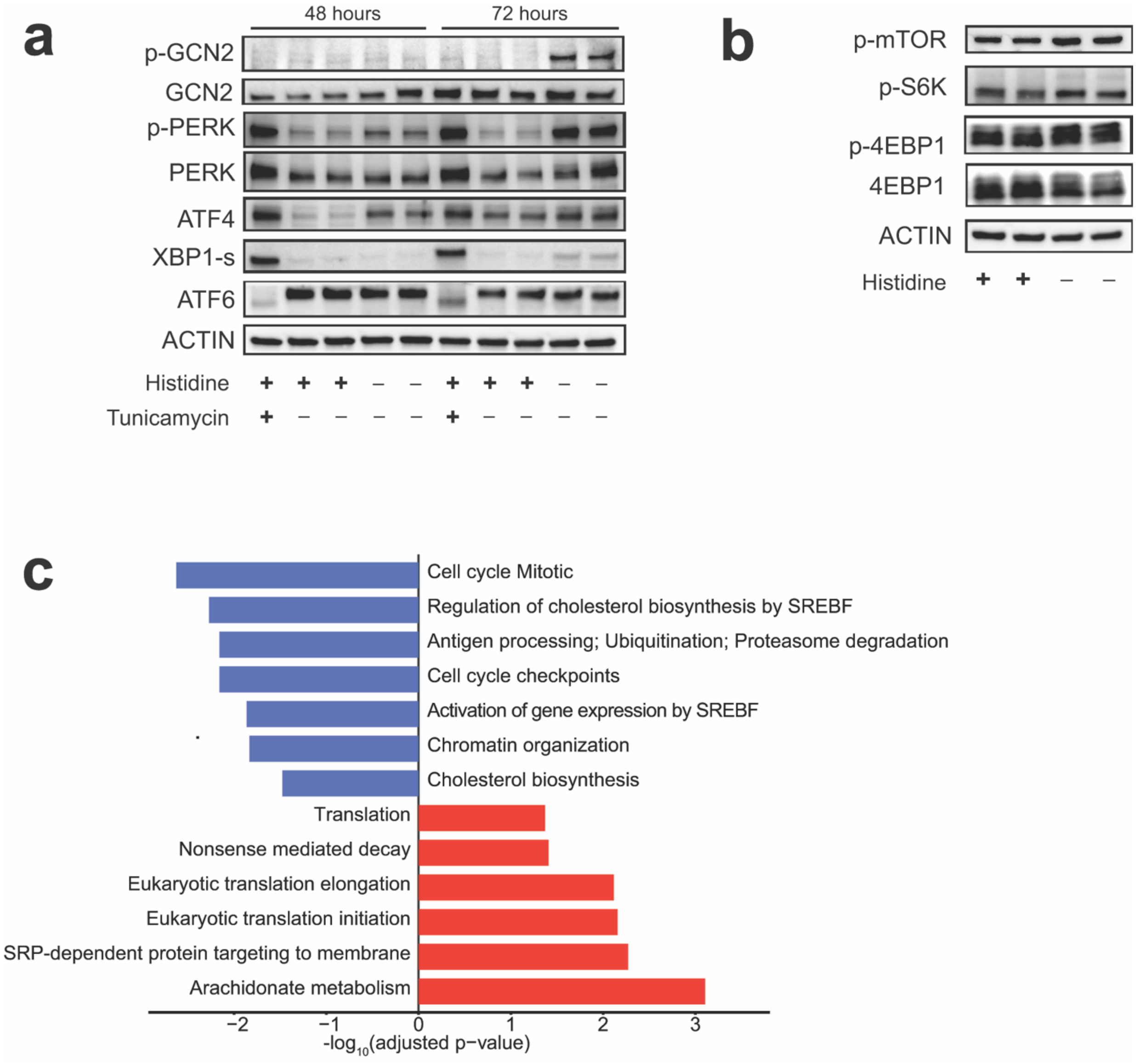


**Extended Data Fig. 8 | Stress response pathway activation and proteome-wide responses to histidine depletion. a**, Western blot analyses of phospho-GCN2, total GCN2, phospho-PERK, total PERK, ATF4, SREBF2 and ACTIN in Jurkat cells grown in 25% histidine for 48 or 72 h and treated with either DMSO or tunicamycin to induce the unfolded protein response. **b**, Western blot analysis of mTOR signaling pathway components in Jurkat cells cultured in histidine-replete (+) or histidine-depleted (25% His; –) media. Phosphorylation of mTOR, S6K, and 4EBP1 increases upon histidine deprivation, indicating enhanced mTORC1 activity. **c**, Pathway enrichment analysis of differentially expressed proteins in Jurkat T-ALL cells in histidine-depleted (25% His) versus control conditions. Upregulated pathways (red) include translation pathways and protein trafficking, whereas downregulated pathways (blue) involve cholesterol biosynthesis and cell cycle progression.

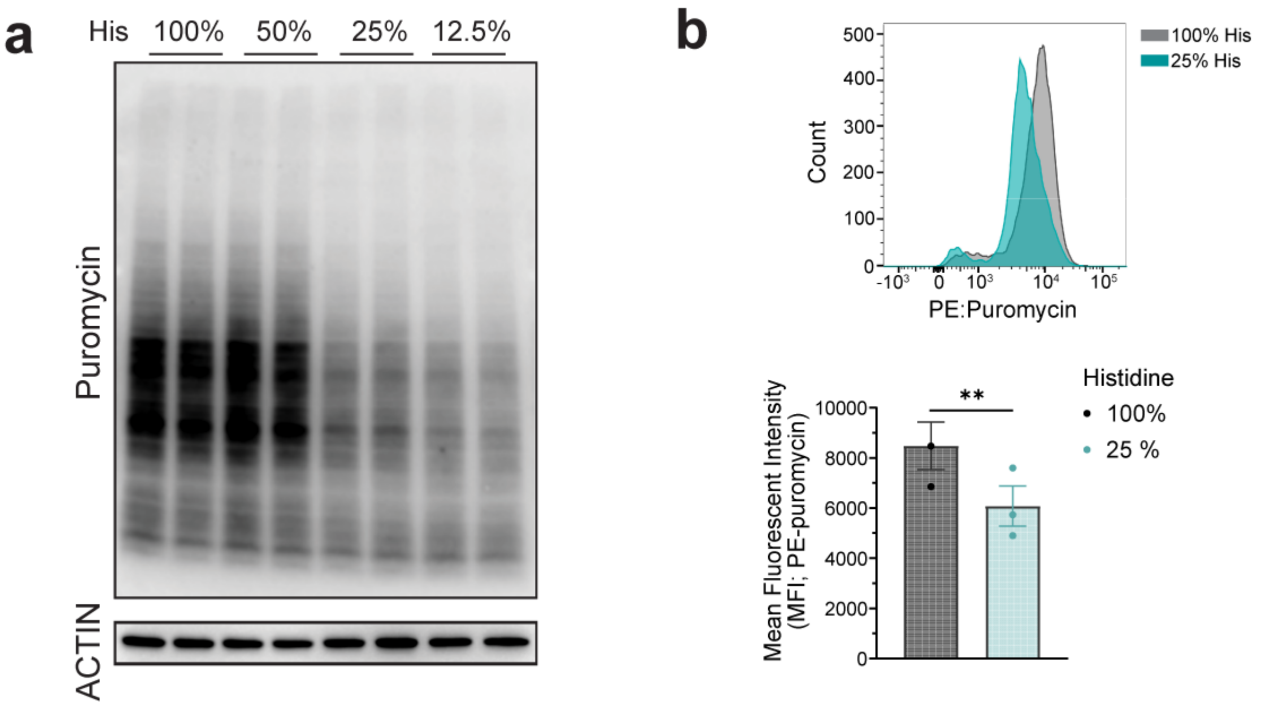


**Extended Data Fig. 9 | Quantitative assessment of translation rates upon histidine depletion. a**, Semi-quantitative analysis of global translation by western blot detection of puromycin-labeled nascent peptides using an anti-puromycin antibody. Stepwise reduction of histidine availability results in a dose-dependent decrease in overall protein synthesis. Representative of three independent experiments. **b**, Flow cytometry-based quantification of puromycin incorporation in Jurkat cells confirms reduced translational activity under histidine-depleted conditions. Mean fluorescence intensity (MFI) of PE–puromycin staining in live cells reflects global protein synthesis rates. Data represent mean ± s.e.m.; $*P < 0.05$, $**P < 0.01$ (two-tailed t-test, n = 4 per group).

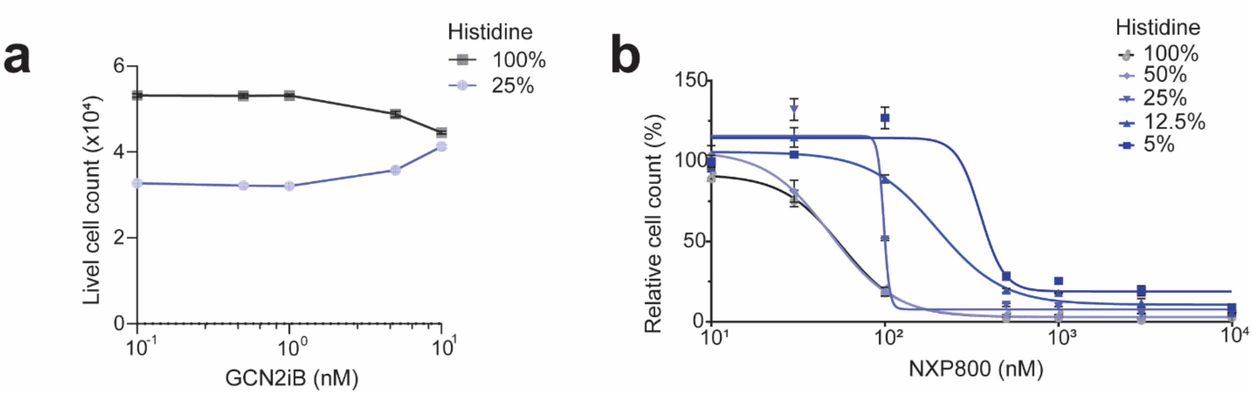


**Extended Data Fig. 10 │ Effects of GCN2 pharmacological inhibition/activation and histidine depletion on the proliferation of Jurkat cells. a,** Drug response profiling with GCN2 inhibitor (GCN2iB) shows rescue of proliferation upon GCN2 inhibition in histidine depleted media. Data are mean ± sem (n=8). **b**, Drug response profiling with GCN2 activator (NXP800) shows reduced sensitivity to GCN2 activation with decreasing levels of histidine. Data are mean ± sem (n=3).

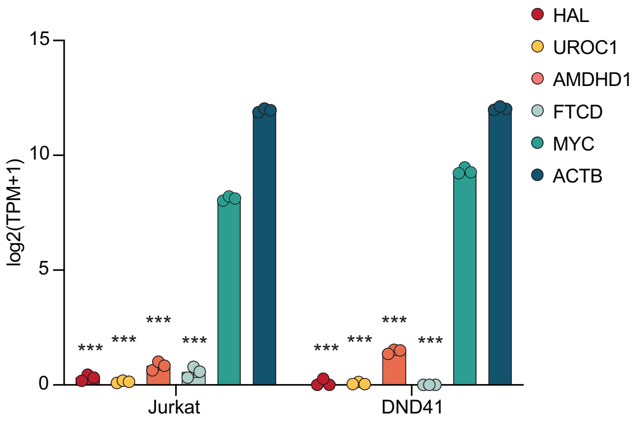


**Extended Data Fig. 11 │ Low expression of histidine catabolism enzymes in T-ALL cells.** TPM counts of Histidine ammonia lyase (HAL), Urocanate (UROC1), Imidazolone-propionate amidohydrolase (AMDHD1), Formimidoyltransferase cyclodeaminase (FTCD), as compared to MYC and b-actin (ACTB) in Jurkat and DND41 cells. Data are mean ± sem; **P* < 0.05; ***P* < 0.01; ****P* < 0.005 (Dunnett's multiple comparison test, n=3 per group).
